# A complex of MAST1 and 14-3-3η regulates Tau phosphorylation in the developing cortex

**DOI:** 10.1101/2025.07.09.663707

**Authors:** Sumire Antonioli, Patrick Heisterkamp, Weiqiang Chen, Dorothea Anrather, Markus Hartl, Maria-Fernanda Martinez-Reza, Ratna Tripathy, Michael Schutzbier, Karl Mechtler, David A. Keays, Thomas A. Leonard

## Abstract

The MAST family of serine/threonine kinases has been implicated in a spectrum of human neurodevelopmental disorders. However, little is known about their biological function or regulation. Seeking to fill these gaps in our knowledge, we have identified upstream and downstream partners of MAST1. 14-3-3η, a neuronal 14-3-3 paralog, specifically interacts with MAST1 at two regulatory serines, S90 and S161. PAK, a neuronal regulator of the actin cytoskeleton, phosphorylates MAST1 to regulate its interaction with 14-3-3η. Exploiting mouse models of human Mega Corpus Callosum Syndrome (MCC) and whole brain phosphoproteomics, we identify the microtubule-associated protein Tau as a substrate of MAST1. We show that pathogenic MAST1 mutations perturb protein function either through misfolding or attenuation of kinase activity. Our data is consistent with a model in which the MAST kinases couple PAK, a neuronal regulator of the actin cytoskeleton, to microtubule remodeling during the differentiation and specification of cortical neurons.

## Introduction

The differentiation of cortical neurons is an essential and tightly regulated process, dysregulation of which is associated with a variety of neurodevelopmental disorders (NDDs). A rare NDD called Mega-Corpus-Callosum Syndrome with Cerebellar Hypoplasia and Cortical Malformations (MCC-CH-CM) is characterized by enlargement of the corpus callosum, epilepsy, and severe cognitive and motor impairment ^1^. Whole exome sequencing of a cohort of seven patients revealed that six harbored *de novo* mutations in a gene called Microtubule Associated Serine/Threonine Kinase 1 (MAST1). A mouse model that recapitulates a heterozygous trinucleotide deletion (L278del) identified in an MCC-CH-CM patient mirrors multiple aspects of the disease, including the enlarged corpus callosum. The identification of additional MCC patients bearing mutations in MAST1, as well as *de novo* mutations in MAST1, MAST3, and MAST4 associated with epilepsy, autism, cerebral palsy, and intellectual disability ^2–9^ has provided further evidence for the involvement of the MAST kinases in neurodevelopment. Despite their association with human NDDs, very little is known about the biological function, structure, or regulation of any MAST kinase.

The MAST kinases are large, ∼200 kDa proteins that first arose ∼600 million years ago in Cnidaria and Ctenophora, the first organisms with a primitive nervous system ^10^. They contain three domains; an N-terminal domain of unknown function (DUF) which harbors a four-helix bundle (4HB), a catalytic Ser/Thr kinase domain, and a C-terminal PDZ domain. The kinase domain belongs to the AGC family of protein kinases, characterized by the presence of a regulatory C-terminal tail ^11^. Although many eukaryotic protein kinases are regulated by activation loop phosphorylation ^12^, the MAST kinases possess a glutamine in place of the canonical serine or threonine.

MAST1 is expressed in post mitotic cortical neurons throughout neurodevelopment ^1,13,14^, associating with the microtubule cytoskeleton via unknown microtubule associated proteins (MAPs) ^1^. It has been reported that the PDZ domain of MAST1 interacts with the adaptor protein β2-syntrophin, bridging it to the dystrophin/utrophin network at postsynaptic densities ^13^. Little is known about MAST substrates. MAST1/2/3 and KIN-4 (the sole homolog of MAST kinases in *C. elegans*) reportedly phosphorylate the C-terminus of PTEN in a PDZ domain-dependent manner, however, this interaction has not been confirmed *in vivo* ^15–17^. In addition, Mitogen-Activated Protein Kinase Kinase (MEK) ^18^ and cAMP-regulated phosphoprotein of 16 kDa (ARPP-16) ^19^ are also reported substrates. However, these findings are largely based on immunoprecipitated proteins and lack kinase-dead controls, while the substrates neither conform to canonical AGC kinase substrate motifs, nor establish a functional link to development of the corpus callosum.

Here, we identify 14-3-3η as an interaction partner of MAST1. We show that 14-3-3η binds specifically to two conserved phosphorylation sites in MAST1, S90 and S161. We identify p21-activated kinase (PAK) as the upstream kinase that phosphorylates MAST1 on S90, and S161 as a regulatory autophosphorylation site. We demonstrate that MAST1 phosphorylated on either S90 or S161 forms a stable complex with 14-3-3η *in vitro*. Using two mouse models of patient-derived MCC-CH-CM mutations, we identify the microtubule-associated protein Tau as a substrate of MAST1 and demonstrate that MAST1 phosphorylates Tau *in vitro*. Finally, we show that MCC-CH-CM-associated disease mutations perturb MAST1 kinase activity either through misfolding of its 4HB domain or disruption of its catalytic site.

## Results

### MAST1 interacts with 14-3-3 proteins via S90 and S161

To shed light on the function and regulation of the MAST kinases, we first sought to identify MAST1 interactors by affinity-purification mass spectrometry (AP-MS) in HEK293 cells. All MAST1 constructs employed in this study are depicted in Figure 1A. MAST1^DKP^, which includes all three folded domains, was N-terminally tagged with EGFP and overexpressed in HEK293 cells (Figure 1B). Comparison of the EGFP-MAST1-bound proteins with an EGFP-only control revealed a MAST1-dependent 6-to 30-fold enrichment of six 14-3-3 paralogs (η, γ, ε, β, ζ, and θ) (Figure 1C, Supplementary Files 1-2). 14-3-3 proteins are known to bind specifically to phosphorylated serines ^20^. Analysis of MAST1’s amino acid sequence by 14-3-3Pred ^21^ revealed two putative 14-3-3 binding motifs ^22,23^ surrounding S90 and S161. Both sites exhibit near-invariant sequence conservation over 600 million years. To visualize a potential interaction between MAST1 and 14-3-3, we predicted the structure of MAST1^D^ and the seven mammalian 14-3-3 isoforms, with a 2:2 stoichiometry implied by the dimeric nature of 14-3-3 proteins. AlphaFold ^24,25^ generated five near-identical models for each prediction with an RMSD ranging between 0.439 and 4.286, placing the unstructured N-terminal region of MAST1 in the cradle of 14-3-3 (Figure 1D-E; 14-3-3η shown as representative). Superposition of the resulting structure with an experimentally determined structure of 14-3-3η in complex with a bound phospho-peptide ^26^ revealed that either residue S90 (Figure 1D) or S161 (Figure 1E) of MAST1 was consistently placed in the same position as the bound phospho-serine of the ligand.

**Figure 1.**
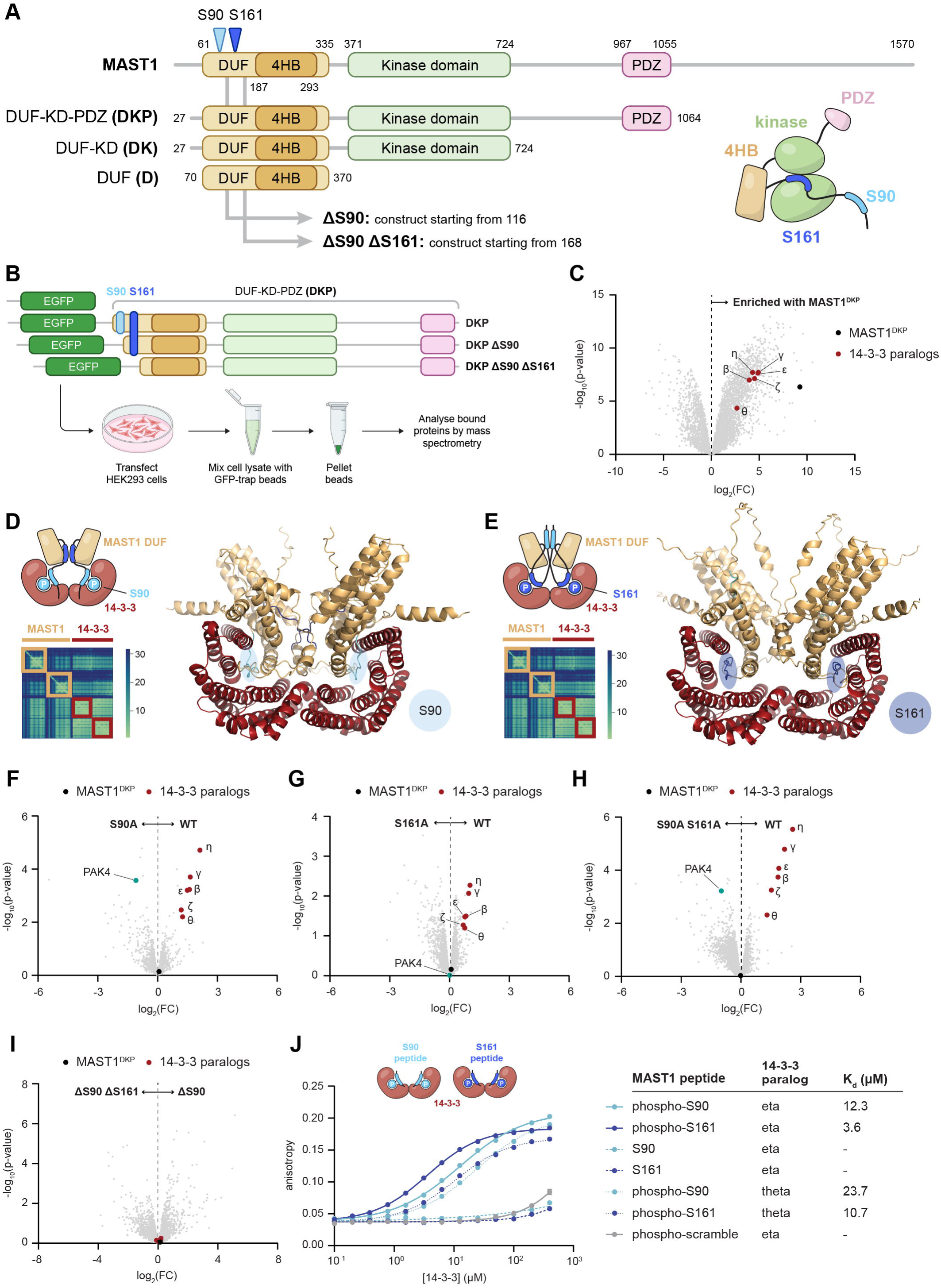
MAST1 interacts with 14-3-3 proteins via S90 and S161. A. Domain architecture of *Mus musculus* MAST1. Construct boundaries and nomenclature for all constructs used in this study are defined. DUF = domain of unknown function, 4HB = four-helix bundle, PDZ = postsynaptic density protein (PDS95) Drosophila disc large tumor suppressor (DlgA) and Zonula occludens-1 protein (zo-1). Right: schematic representation of MAST1 to be employed throughout the figures. B. Construct boundaries and experimental setup for EGFP-MAST1^DKP^ AP-MS in HEK293 cells. C. Volcano plot displaying the enriched interactome of EGFP-MAST1^DKP^ (right) vs EGFP (left) from AP-MS, *n* = 3. D. AlphaFold3 ^25^ model and pair alignment error (PAE) plot of 14-3-3η dimer (red) and MAST1^D^ dimer (gold), when region spanning MAST1 S90 (light blue bubble) interacts with 14-3-3 cradle. E. AlphaFold3 ^25^ model and PAE plot of 14-3-3η dimer (red) and MAST1^D^ dimer (gold), when region spanning MAST1 S161 (dark blue bubble) interacts with 14-3-3 cradle. F. Volcano plot displaying the enriched interactome of EGFP-MAST1^DKP^ WT (right) vs EGFP-MAST1^DKP^ S90A (left) from AP-MS, *n* = 3. G. Volcano plot displaying the enriched interactome of EGFP-MAST1^DKP^ WT (right) vs EGFP-MAST1^DKP^ S161A (left) from AP-MS, *n* = 3. H. Volcano plot displaying the enriched interactome of EGFP-MAST1^DKP^ WT (right) vs EGFP-MAST1^DKP^ S90A S161A (left) from AP-MS, *n* = 3. I. Volcano plot displaying the enriched interactome of EGFP-MAST1^DKP^ ΔS90 (right) vs EGFP-MAST1^DKP^ ΔS90 ΔS161 (left) from AP-MS, *n* = 3. J. Binding curves for 14-3-3η or θ and MAST1 peptides spanning S90 or S161, measured by fluorescence anisotropy. Data are presented as mean values ± S.E., *n* = 3. Data were fit with a one-site binding model and the K_d_ derived from curve fitting.

To probe whether the N-terminal region of MAST1 binds to 14-3-3, we compared truncation constructs of MAST1 lacking S90, or both S90 and S161 (Figure 1B). By performing pairwise comparison of the interactomes of the truncation constructs, we observed significant enrichment of six 14-3-3 paralogs by MAST1^DKP^ when compared to either MAST1^DKP^ ΔS90 (Figure S1A) or MAST1^DKP^ ΔS90 ΔS161 (Figure S1B). To determine the site specificity of the interaction, we next performed AP-MS comparisons between MAST1^DKP^ WT and S90A or S161A point mutants. Mutation of either S90 or S161 resulted in a significant loss of 14-3-3 binding to MAST1 (Figure 1F-G), while double mutation of both residues resulted in an apparently synergistic loss of 14-3-3 proteins (Figure 1H). The absence of 14-3-3 enrichment in MAST1^DKP^ ΔS90, which includes S161 but not S90 (Figure 1I), suggests that 14-3-3 binding to S161 is dependent on 14-3-3 first binding to S90. 14-3-3η was reproducibly the most highly enriched paralog of the six, despite having the lowest abundance in HEK293 cells ^27^ (Figure S1C).

We further confirmed the interaction between purified 14-3-3η to MAST1 by determining its binding affinity to phosphorylated peptides corresponding to the S90 and S161 motifs *in vitro* (Figure 1J). We confirmed the correct mass of all recombinant proteins and derivatives thereof employed in this study by mass spectrometry (Figure S2). Consistent with the AP-MS results, 14-3-3η bound to both phospho-S90 (p-S90) and phospho-S161 (p-S161) peptides, but neither to their unphosphorylated counterparts nor a phosphopeptide with a scrambled amino acid sequence. 14-3-3θ, which exhibited the lowest enrichment by AP-MS, bound MAST1 phosphopeptides with a correspondingly lower affinity.

Finally, single cell sequencing has shown that MAST1 and 14-3-3η are expressed simultaneously in developing cortical neurons in mice ^28^ (Figure S3A-D). In summary, 14-3-3η binds to MAST1 phosphorylated on either S90 or S161.

### PAK4 phosphorylates MAST1 on S90

The phosphorylation-dependent interaction of MAST1 with 14-3-3η immediately begs the question of which kinase(s) phosphorylate S90 and S161. To identify candidate upstream kinases for the conserved S90 site (Figure 2A), we employed the ScoreSite prediction algorithm ^29^, which identified PAK4-6 as the highest ranked kinases (Figure 2B). Curiously, we observed significant enrichment of PAK4 in all AP-MS experiments involving MAST1 S90A (Figure 1F, 1H), suggesting that endogenous PAK4 may have been trapped on a non-phosphorylatable substrate. While PAK4 is relatively abundant in HEK293 cells (280 nM), PAK5 and PAK6 are not detected ^30^ (Figure S4A). PAK4-6 are group II PAKs with a consensus motif that differs from group I PAKs (PAK1-3) at positions P+2 and P+3 (Figure S4B). PAK4-6 recognize an alanine at the P+2 position, in contrast to the tyrosine or phospho-tyrosine favored by PAK1-3, and PAK4 has a strong preference for a serine at the P+3 position ^31^. The S90 motif of MAST1 encodes alanine and serine in the P+2 and P+3 positions respectively (Figure 2A). Consistent with PAK4 being the upstream kinase of MAST1 in HEK293 cells, we observed that purified, recombinant PAK4 phosphorylates MAST1^D^ in an *in vitro* radiometric kinase assay while a kinase-dead mutant did not (Figure 2C). Mass spectrometry analysis revealed that S90 was phosphorylated with the highest spectral count (Figure 2D-E, Supplementary File 3). We also observed S77 to be phosphorylated, likely due to the similarity in the surrounding amino acid sequence. Single cell RNA sequencing data of the developing mouse brain shows a significant overlap in the transcripts of MAST1 and PAK6, whereas PAK4 is barely detected and PAK5 is not detected at all (Figure S4C-H). These findings imply that a group II PAK is an upstream regulator of MAST1 signaling in the developing cortex, with PAK6 being the strongest candidate.

**Figure 2.**
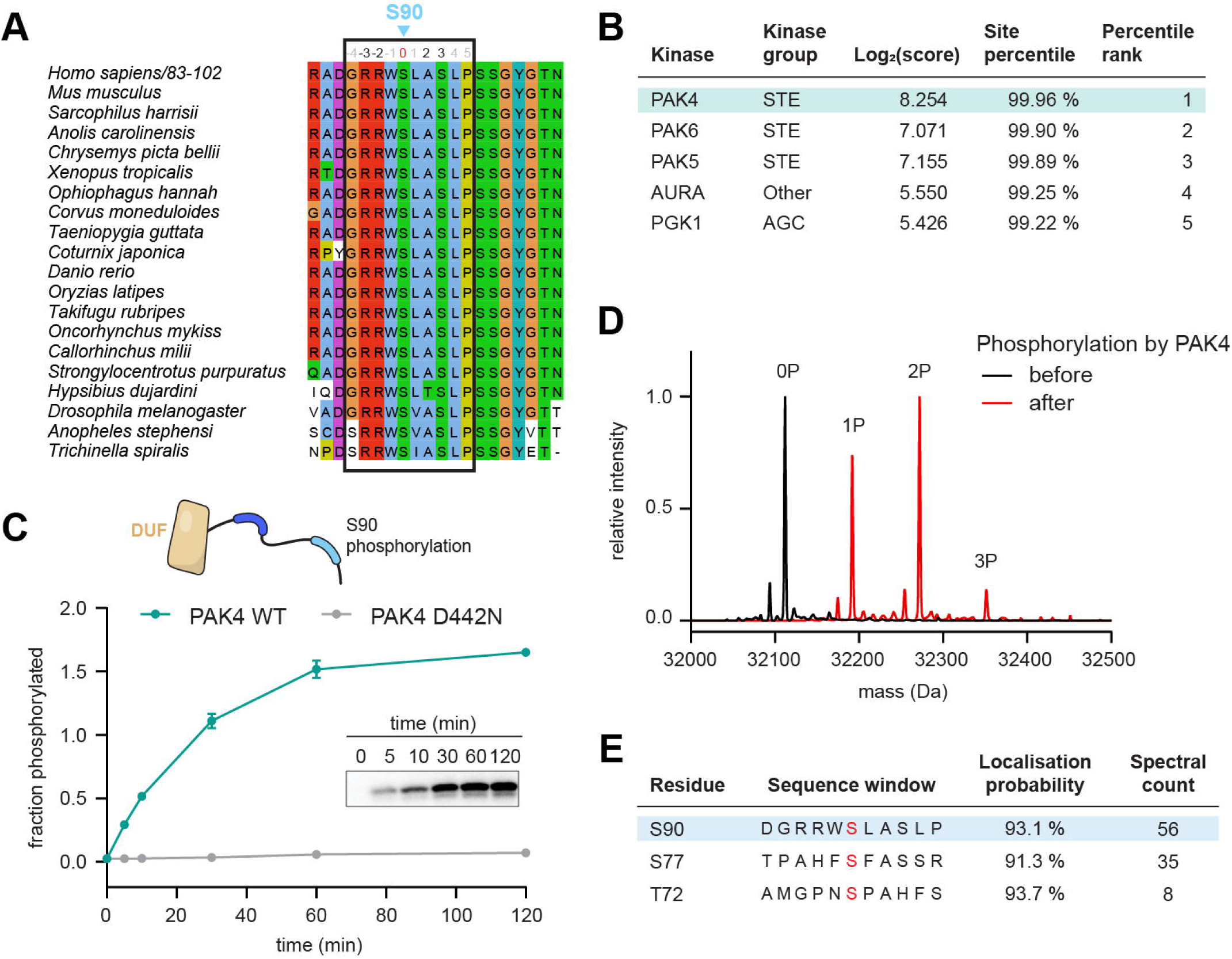
PAK4 phosphorylates MAST1 on S90. A. Multiple sequence alignment of MAST1 residues surrounding S90 from *Nematoda* to *Mammalia*. S90 residue highlighted above, sequence used for ScoreSite prediction outlined in black box. B. ScoreSite ^29^ prediction results using input sequence GRRWs*LASLP (s* represents S90 phosphosite). Top five hits shown in table, PAK4 highlighted in turquoise. C. Radiometric kinase assay by PAK4 WT or PAK4 kinase dead (D442N) on MAST1^D^. Data are presented as mean values ± S.E., *n* = 3. Representative phosphorimage shown in inset. D. Intact MS analysis of MAST1^D^ before (black) and after (red) phosphorylation by PAK4 WT shown in panel C. Intensity of both samples were normalized to a scale of 0 to 1. Addition of phosphate (1P, 2P etc.) detected by each 80 Da shift. E. Tandem MS analysis of MAST1^D^ after phosphorylation by PAK4 WT shown in panel C. Phosphorylated serines are indicated in red. Top three sites (by spectral count) shown in table, S90 highlighted in light blue. See Supplemental File 3 for full data.

### MAST1 S161 is an autoregulatory motif

Having identified the upstream kinase for S90, we next sought to identify the kinase that phosphorylates S161. Inspection of the conserved motif surrounding S161 (Figure 3A) revealed the presence of a canonical AGC kinase consensus sequence, characterized by arginine in the P-5 and P-3 positions, as well as a large hydrophobic residue in the P+1 position (Figure 3B) ^11^. This motif is near invariant in all MAST kinases (Figure 3C), implying an evolutionarily conserved function. Since MAST kinases are, themselves, members of the AGC family, this observation suggested that S161 may represent an autoregulatory phosphorylation site. AlphaFold predictions of a complex between the MAST1 kinase domain and the DUF reproducibly converged on a model that places S161 in the substrate-binding cleft of the kinase domain, canonically poised for phospho-transfer (Figure 3D, S5A). In an *in vitro* kinase assay, recombinant MAST1^DK^ ΔS90 (Figure S5B-C) incubated with ATP was able to autophosphorylate, while a kinase-dead MAST1 was completely inactive (Figure 3E-F). To determine the site-specificity of the reaction, we performed tandem mass spectrometry, which revealed phosphorylation of S161 (Figure 3G, Supplementary File 3). However, the reaction was inefficient and resulted in multiple phosphorylated sites, suggesting that unknown factors may be required for full and specific MAST1 activity. Finally, we measured the binding affinity of MAST1^DK^ ΔS90 ΔS161 (Figure S5D-E) for a peptide corresponding to the S161 motif (Figure 3H). A peptide bearing the native sequence of MAST1 bound to the kinase with a low affinity consistent with the known strength of kinase-substrate interactions ^32^. Mutation of the arginines at P-3 and P-5 reduced the binding affinity, which was further reduced by mutation of the leucine in the P+1 position (Figure 3H). These findings are consistent with a specific interaction of the conserved S161 motif with the MAST1 kinase domain. In summary, MAST1 undergoes autophosphorylation at S161, but the factors which control the rate and specificity of the reaction are unknown.

**Figure 3.**
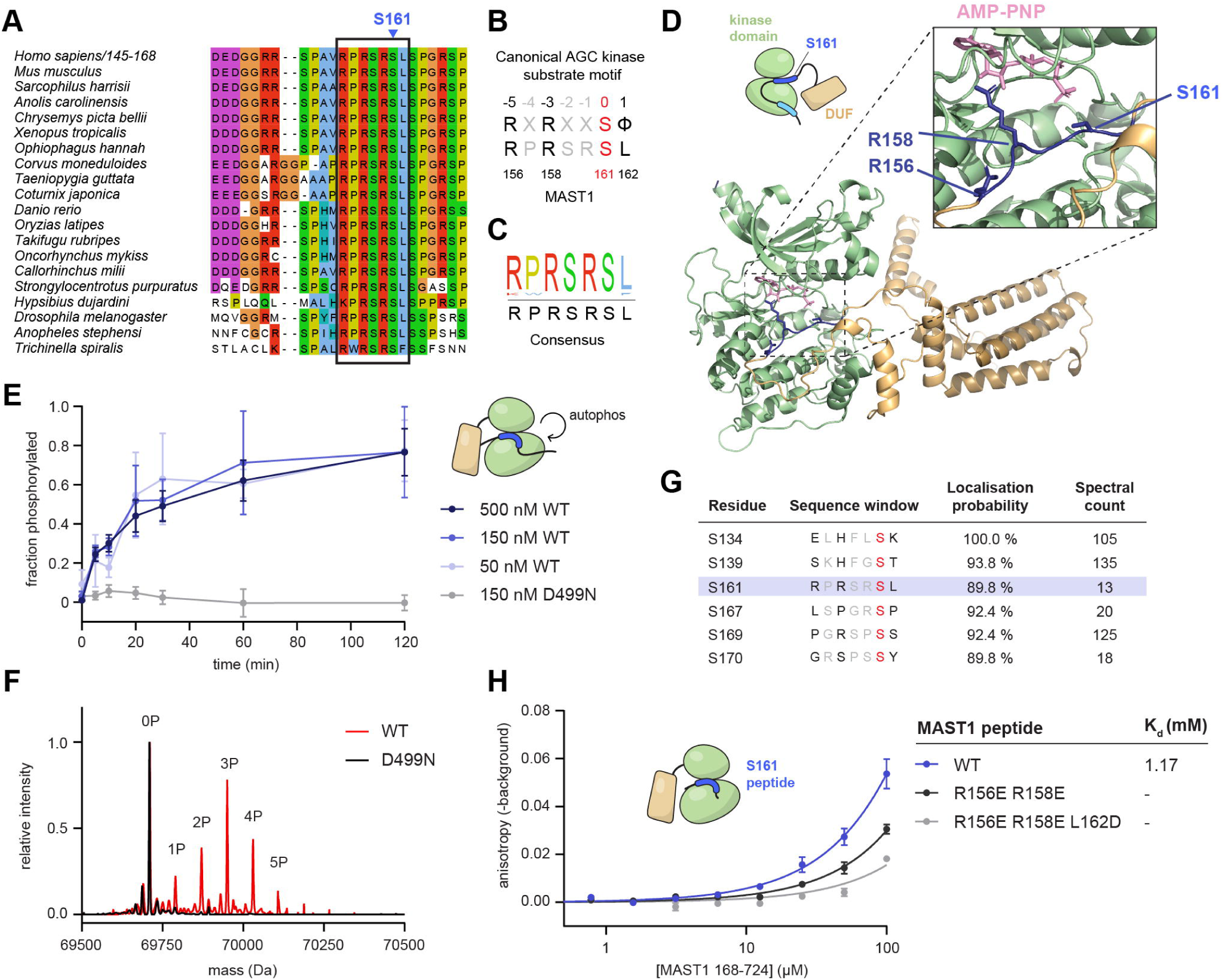
MAST1 S161 is an autoregulatory motif. A. Multiple sequence alignment of MAST1 residues surrounding S161 from *Nematoda* to *Mammalia*. S161 residue highlighted above, AGC kinase recognition region outlined in black box. B. Alignment of canonical AGC kinase substrate motif with MAST1^156–162^, where S161 is the phosphorylatable residue (red) and R156 (-5), R158 (-3), and L162 (+1) are in positions specifically involved in substrate recognition (black). C. Consensus sequence of MAST1^156–162^ from sequence alignment in panel A. D. AlphaFold2 ^24^ model of MAST1 DUF (gold) and kinase domain (green), with region spanning S161 (dark blue) positioned in the catalytic site of MAST1. AMP-PNP molecule (pink) was incorporated into the model by superposition of MAST1 kinase domain with X-ray structure of Akt (PDB ID 1O6K) ^77^. E. Radiometric autophosphorylation assay with various concentrations of MAST1^DK^ ΔS90 WT or kinase dead (D499N). Data are presented as mean values ± S.E., *n* = 3. F. Intact MS analysis of MAST1^DK^ ΔS90 WT (red) and kinase dead (D499N) (black) after autophosphorylation shown in panel E. Intensity of both samples were normalized to a scale of 0 to 1. Addition of phosphate in WT sample (1P, 2P etc.) detected by each 80 Da shift. G. Tandem MS analysis of MAST1^DK^ ΔS90 WT after autophosphorylation shown in panel E. Phosphorylated serines are indicated in red and residues involved in AGC kinase binding (-5, -3, +1) are in black. Top six serine sites (by spectral count) shown in table, S161 highlighted in dark blue. See Supplemental File 3 for full data. H. Binding curves for MAST1^DK^ ΔS90 ΔS161 and MAST1 peptides spanning S161, measured by fluorescence anisotropy. Data are presented as mean values ± S.E., *n* = 3. Data were fit with a one-site binding model and the K_d_ derived from curve fitting.

### MAST1 phosphorylated on S90 or S161 forms a complex with 14-3-3η *in vitro*

Having established that MAST1 interacts specifically with 14-3-3η in a phosphorylation-dependent manner both in cells and *in vitro*, we sought to reconstitute stable complexes of the purified proteins *in vitro*. To first reconstitute S90-phosphorylated MAST1:14-3-3η, we co-expressed MAST1^DK^ with 14-3-3η and PAK4^FL^ in baculovirus-infected Sf9 insect cells. Affinity purification of the GST-tagged 14-3-3η yielded a complex with MAST1 which was stable by size-exclusion chromatography (SEC) (Figure 4A-B). Unexpectedly, however, MAST1 was found to be phosphorylated not only on S90 but also on S161 and S163 by tandem mass spectrometry, with higher spectral counts for p-S161 and p-S163 than p-S90 (Figure 4C, Supplementary File 3). The origin and significance of MAST1 phosphorylation on S163 in the insect cells is unclear, but a phospho-peptide corresponding to the S163 motif is incapable of binding to 14-3-3η (Figure S6A). Therefore, the complex of MAST1 and 14-3-3η isolated from insect cells co-expressing PAK4 is mediated by phosphorylation of S90, S161, or a combination of both. This suggests that PAK4-mediated MAST1 phosphorylation on S90 promotes the acquisition of p-S161, presumably by autophosphorylation, but this will require further investigation. To determine the stoichiometry of the complex, we measured the mass of the particles by mass photometry ^33^. Unfortunately, at the concentrations required for mass photometry, the complex dissociated into its constituent components (Figure S6B). We therefore employed glutaraldehyde cross-linking to stabilize the complex (Figure S6C) and measure its stoichiometry. Particles of MAST1 and 14-3-3η were observed to have an average mass of 132 kDa, which corresponds to a 1:2 complex (Figure 4D, Figure S6D-F). The 1:2 stoichiometry was further supported by gel densitometry analysis of the complex in Figure 4B.

**Figure 4.**
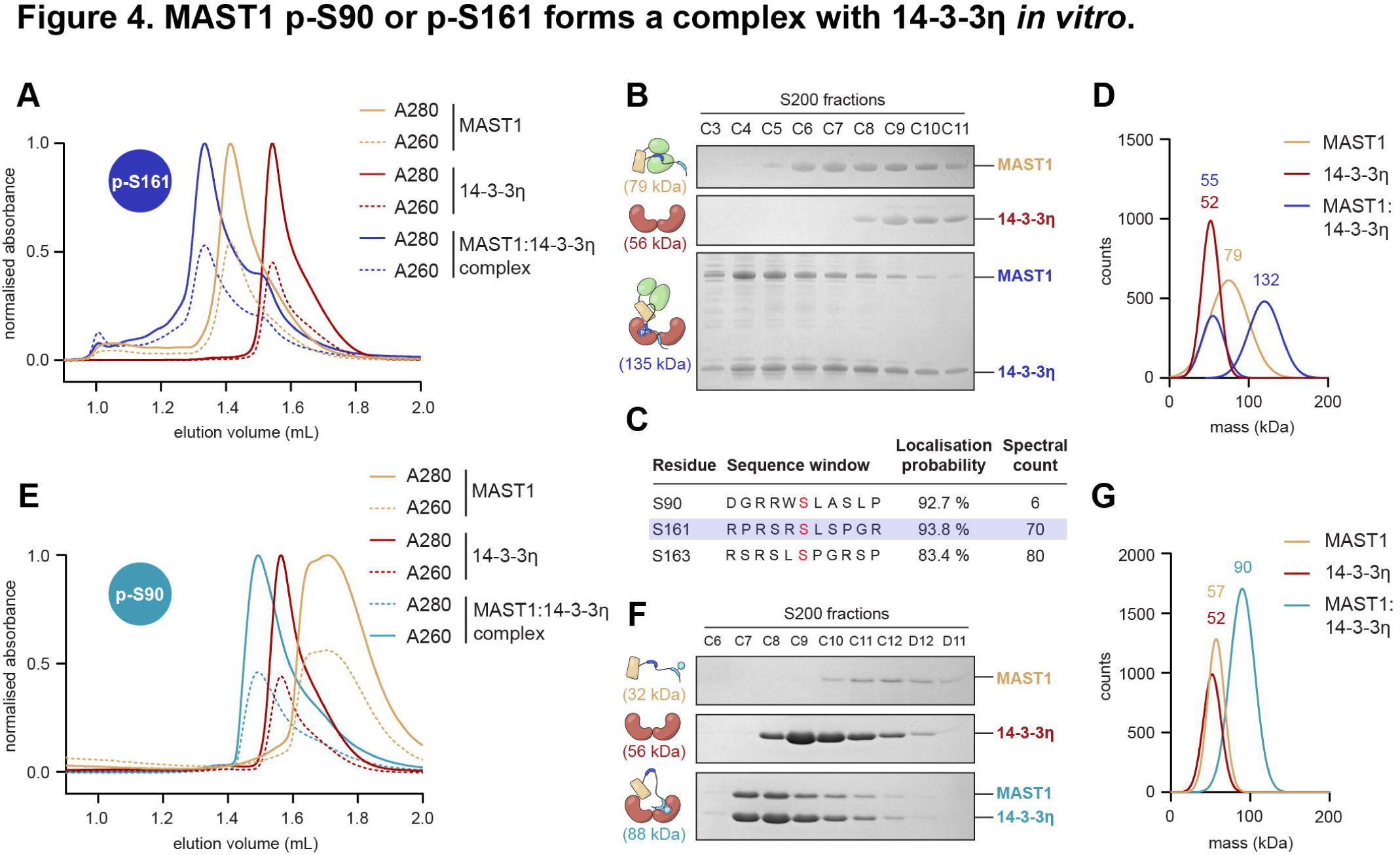
MAST1 p-S90 or p-S161 forms a complex with 14-3-3η *in vitro*. A. Analytical size exclusion chromatography (SEC) elution profiles of purified MAST1^DK^ (yellow), purified 14-3-3η (red), and a co-purified complex of MAST1^DK^ and 14-3-3η (dark blue). B. Coomassie-stained SDS-PAGE gel of SEC fractions spanning elution peaks in panel A. Theoretical masses of each protein/complex shown in parentheses. C. Tandem MS analysis of co-purified MAST1^DK^ and 14-3-3η complex to determine phosphorylation state of MAST1 in the complex. Phosphorylated serines are indicated in red. S90, S161 (highlighted in dark blue), and S163 sites shown in table. See Supplemental File 3 for full data. D. Gaussian fits from mass photometry of purified MAST1^DK^ (yellow), purified 14-3-3η (red), and a co-purified complex of MAST1^DK^ and 14-3-3η (dark blue). Experimentally measured masses of each protein/complex indicated above the Gaussian fit. E. Analytical SEC elution profiles of purified MAST1^D^ p-S90 (yellow), purified 14-3-3η (red), and an *in vitro*-formed complex of MAST1^D^ p-S90 and 14-3-3η (light blue). F. Coomassie-stained SDS-PAGE gel of SEC fractions spanning elution peaks in panel E. Theoretical masses of each protein/complex shown in parentheses. G. Gaussian fits from mass photometry of purified MAST1^D^ p-S90 (yellow), purified 14-3-3η (red), and an *in vitro*-formed complex of MAST1^D^ p-S90 and 14-3-3η (light blue). Experimentally measured masses of each protein/complex indicated above the Gaussian fit.

Since efforts to obtain S90-phosphorylated MAST1 in complex with 14-3-3η by co-expression with PAK4 yielded complexes with a mixture of S90- and S161-phosphorylated MAST1, we reconstituted the S90-phosphorylated MAST1:14-3-3η complex *in vitro* to assess its stoichiometry. Purified MAST1^D^, phosphorylated with PAK4^FL^ and subsequently incubated with 14-3-3η at a 1:1 stoichiometry, was purified by analytical SEC (Figure 4E). Complex formation was evidenced by the increase in hydrodynamic radius of MAST1 and the co-elution of the two proteins. Gel densitometry of the co-eluting peak again indicated a 1:2 complex (Figure 4F). This was confirmed by glutaraldehyde cross-linking mass photometry, where particles of S90-phosphorylated MAST1 crosslinked to 14-3-3η were observed to have an average mass of 90 kDa, corresponding to a 1:2 complex (Figure 4G, Figure S6D, G-H). In summary, we could reconstitute MAST1:14-3-3η complexes *in vitro*, mediated by either S90 and S161 phosphorylation, and both assemble with a stoichiometry of 1:2.

### MAST1 G519S mice exhibit an enlarged corpus callosum

To investigate the mechanisms of MCC-CH-CM pathogenicity, we next focused on the molecular consequences of patient-derived MAST1 mutations. MCC-CH-CM patients with heterozygous mutations in MAST1 exhibit gross morphological alterations in their brain structure, with a significantly enlarged corpus callosum ^1^. Curiously, MAST1 knock-out mice are phenotypically normal ^1^, implying that the disease is driven by a dominant negative effect of the mutant MAST1 and not by haploinsufficiency. Three of these mutations are triplet deletions in the coding sequence of the 4HB domain and one is a missense mutation in the activation loop of the kinase domain that replaces an invariant glycine in the DFG motif with a serine (Figure 5A). A MAST1^L278del/+^ mouse model has previously been generated and exhibits an enlarged corpus callosum and thinner cortex ^1^. The G517S patient mutation (G519S in mice) has, as yet, not been phenotypically characterized in a mouse model. Using CRISPR-Cas9 genome editing, we generated MAST1^G519S/+^ and MAST1^G519S/G519S^ mice. Following pronuclear injection with guides targeting exon 14 (Supplementary Table 1), founder animals were identified by Sanger sequencing (Figure 5B-C) and backcrossed for 10 generations to C57Bl/6J. G519S homozygotes died shortly after birth, whereas heterozygotes survived into adulthood. Analysis of adult G519S/+ mouse brains by Nissl staining confirmed the presence of an enlarged corpus callosum with a reduction in cortical thickness (Figure 5D-F) consistent with the disease state and the L278del mouse model. It is worth noting that while the phenotypes of the G519S/+ mouse and the previously described L278del/+ mouse are qualitatively similar, the L278del/+ line displays a more pronounced enlargement of the corpus callosum, hinting towards a more severe molecular disruption caused by the L278del mutation. Analysis of MAST protein levels in P0 cortical samples revealed a dose dependent decrease in the levels of MAST1, MAST2 and MAST3 in the G519S/+ mouse compared to +/+ (Figure 5G-H) consistent with a dominant negative mechanism. This implies that the disease phenotype stems from a perturbation of MAST signaling in the developing brain.

**Figure 5.**
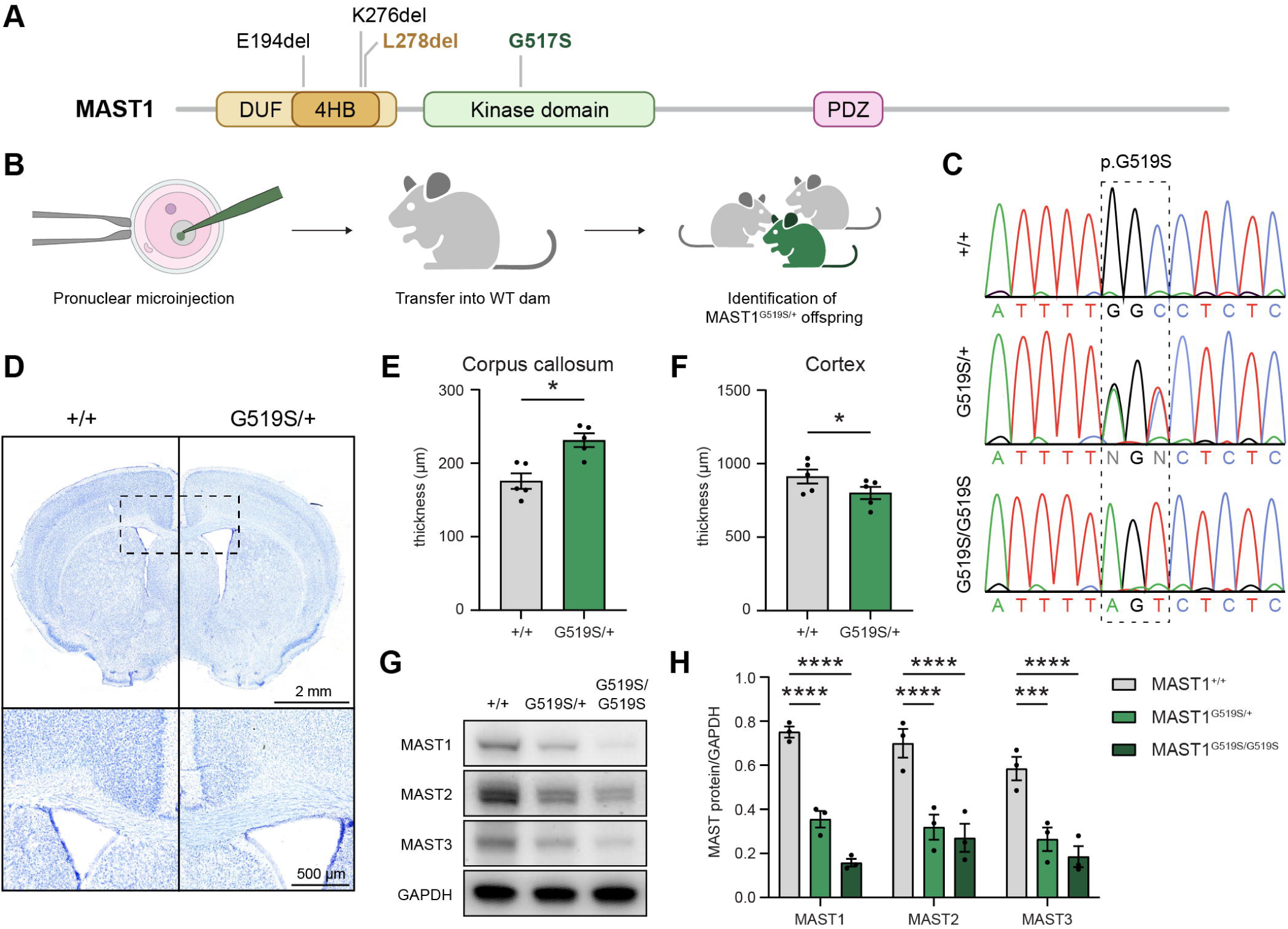
MAST1 G519S mice exhibit an enlarged corpus callosum. A. Schematic of MCC-CH-CM patient-derived mutations in MAST1 protein. G519S (mouse numbering) corresponds to G517S in humans. Mouse models have been generated for the highlighted L278del and G519S variants. B. G519S animals were generated by injection of a single strand DNA repair template and Cas9/guide RNA into the pronucleus of mouse zygotes. Founder animals were identified by Sanger sequencing and backcrossed to C57Bl/6J animals for 10 generations. C. Sequencing traces of MAST1 allele for control (+/+), heterozygous (G519S/+), and homozygous animals (G519S/G519S). D. Nissl stain of coronal 8-week-old control (+/+) and heterozygous brains (G519S/+) shows an enlargement of the corpus callosum in mutants. Bregma =0.86 mm. E. Quantification of corpus callosal thickness in control (+/+) and heterozygous brains (G519S/+) (*n* = 5, P < 0.05, paired, two-sided t-test with Bonferroni correction). Data are presented as mean values ± S.E. F. Quantification of cortical thickness in control (+/+) and heterozygous brains (G519S/+) (*n* = 5, P < 0.05, paired, two-sided t-test with Bonferroni correction). Data are presented as mean values ± S.E. G. Western blot analysis on P0 cortical lysates for MAST1, MAST2, and MAST3 in control (+/+), heterozygous (G519S/+) and homozygous (G519S/G519S) animals. H. Quantification of western blot in panel F (*n* = 3, * p < 0.05; ** p < 0.01; ***, p < 0.001; **** p < 0.0001, two-way ANOVA with Dunnet’s multiple comparisons test). Intensity of bands for MAST family members was normalized to corresponding GAPDH band. Data are presented as mean values ± S.E.

### Tau is a MAST1 substrate

To identify candidate substrates of MAST1, we exploited the previously published MAST1^L278del/L278del^ mouse model and the MAST1^G519S/G519S^ model generated in this study. To explore how the mutations alter the phosphoproteome in the developing brain we extracted protein lysates from P0 cortices from both controls and homozygotes for the L278del and G519S lines. This time point was selected because the callosal projection neurons are already specified and are extending axons to the contralateral hemisphere ^35^. Following protein isolation, denaturation, reduction and alkylation, isobaric labels were added and samples were enriched for phosphopeptides using titanium dioxide (TiO2) resin. This resulted in the identification by mass spectrometry of 50601 phosophopeptides in the L278del mouse line and 22192 phosphopeptides in the G519S line, of which 1608 and 610 passed the 5% FDR threshold respectively. A comparative analysis identified 70 phosphopeptides, from 38 proteins, that were observed to be differentially phosphorylated in both L278del and G519S mice when compared to littermate controls (Figure 6A, Supplementary File 4). By filtering for hypo-phosphorylated proteins for which the phosphorylated residue was embedded in an AGC kinase consensus motif, we reduced the list of candidate proteins to four (Figure 6B). Of the four candidates, phosphorylation of microtubule-associated protein Tau (MAPT) on S214 was most notable. Tau is a microtubule-associated protein that, like MAST1, is highly expressed in neurons of the developing mouse brain (Figure S7A-B) and the phosphorylation state of S214 is considered to be a biomarker for Alzheimer’s Disease (AD) ^34,36^. Inspection of the amino acid sequence surrounding S214 revealed that it is near-identical to the S161 autophosphorylation site in MAST1 (Figure 6C). Phosphorylation of Tau on S214 was reduced in both L278del and G519S mice compared to littermate controls (Figure 6D-E).

**Figure 6.**
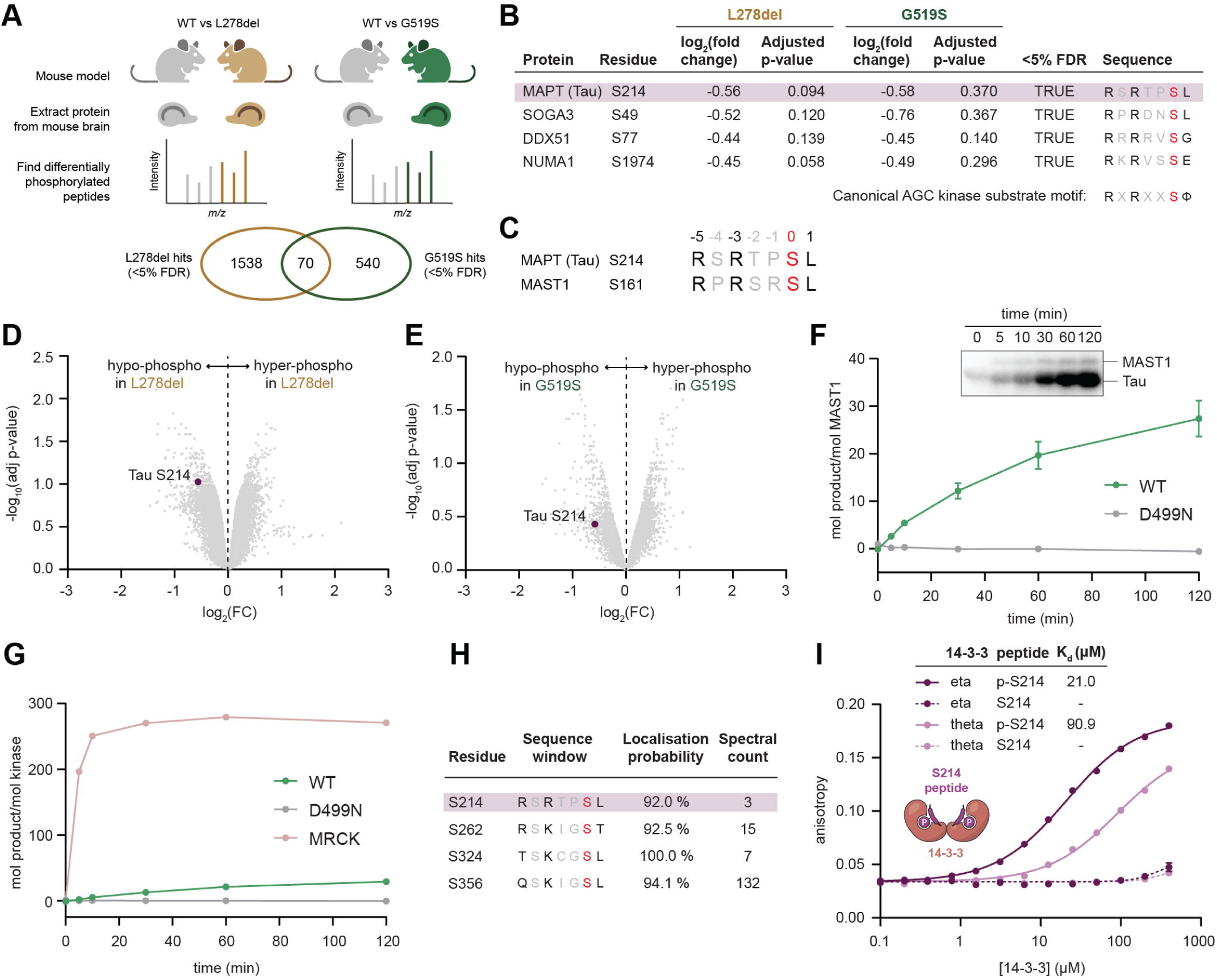
Tau is a MAST1 substrate. A. Schematic of MAST1 WT vs L278del vs G519S phosphoproteomics experiment. Number of significantly differentially phosphorylated peptides obtained in each comparison, and shared in both comparisons, shown in Venn diagram (*n* = 5 WT and homozygous litter-matched P0 mice, LIMMA test with 5% FDR threshold described in Hein *et al* ^30^ was applied). B. Table displaying data from four of the 70 shared phosphosite hits in panel A. The four phosphosites listed all conform to the canonical AGC kinase substrate motif (phosphorylated serines indicated in red). MAPT (Tau) is highlighted in purple. C. Sequence alignment of Tau^209–215^ and MAST1^156–162^, where Tau S214 (substrate) and MAST1 S161 (autophosphorylation site) are aligned. D. Volcano plot of phosphoproteomics analysis showing hyperphosphorylated sites (right) and hypophosphorylated sites (left) for L278del when compared to WT. Tau S214 highlighted in purple. E. Volcano plot of phosphoproteomics analysis showing hyperphosphorylated sites (right) and hypophosphorylated sites (left) for G519S when compared to WT. Tau S214 highlighted in purple. F. Radiometric kinase assay by MAST1^DK^ ΔS90 WT or kinase dead (D499N) on Tau. Data are presented as mean values ± S.E., *n* = 10 for WT, 3 for D499N. Representative phosphorimage shown in inset. G. Radiometric kinase assay by MRCKα on MLC2, overlaid on assay shown in E. H. Tandem MS analysis of Tau after phosphorylation by MAST1^DK^ ΔS90 WT shown in panel E. Phosphorylated serines are indicated in red and residues involved in AGC kinase binding (-5, -3, +1) are in black. S214 site (highlighted in purple) as well as three known Tau phosphorylation sites in its repeat domains shown in table. See Supplemental File 3 for full data. I. Binding curves for 14-3-3η or θ and Tau peptides spanning S214, measured by fluorescence anisotropy. Data are presented as mean values ± S.E., *n* = 3. Data were fit with a one-site binding model and the K_d_ derived from curve fitting.

To assess whether MAST1 can phosphorylate Tau *in vitro*, we performed a kinase assay with purified, recombinant Tau (mouse isoform F). The purity and monodispersity of Tau^FL^ was confirmed by SEC and mass photometry (Figure S7C-D). Wild-type MAST1^DK^ ΔS90 phosphorylated Tau, and a kinase-dead mutant of MAST1 was completely inactive against Tau (Figure 6F). However, the speed of the reaction was lower than what would be expected of a fully active AGC kinase. To benchmark this quantitatively, we compared MAST1 phosphorylation of Tau to the catalytic activity of the closely related AGC kinase Myotonic Dystrophy Kinase-Related Cdc42-binding Kinase α (MRCKα), on its substrate myosin light chain 2 (MLC2). MRCKα exhibited a 50-fold higher catalytic efficiency than MAST1 (Figure 6G). The post-translational modification profile of Tau exhibited multiple phosphorylation sites including S214, as well as three known Tau phosphorylation sites ^37–39^ within its repeat domains (Figure 6H, Supplementary File 3). This phosphorylation pattern is reminiscent of MAST1 autophosphorylation (Figure 3F-G), implying that other factors govern the site-specificity of the reaction. To test whether the catalytic activity of MAST1 is increased when in complex with 14-3-3η, we compared Tau phosphorylation by MAST1^DK^ alone or in complex with 14-3-3η, but observed no significant increase (Figure S7E).

Tau phosphorylated on S214 has previously been reported to bind 14-3-3 proteins ^40,41^. Given the similarity of the S214 motif in Tau to the S161 motif in MAST1 and the observation that MAST1 phosphorylated on S161 interacts with 14-3-3η, this is not surprising. We confirmed that purified 14-3-3η binds a phosphorylated peptide of the Tau S214 motif by fluorescence anisotropy (Figure 6I). As previously observed for p-S161, the binding was phosphorylation-dependent and 14-3-3η exhibited a higher affinity than 14-3-3θ. Taken together, these findings support a model in which MAST1 activity plays a role in the regulation of the microtubule cytoskeleton of neurons, via Tau.

### MCC-CH-CM mutations perturb MAST1 folding and kinase activity

MAST1 proteins in mouse models of patient-derived MCC-CH-CM mutations exert a dominant negative effect by depleting MAST1, 2, and 3 (Figure 5H and Tripathy *et al.* ^1^). To investigate the molecular basis of pathogenicity, we characterized the effect of the four patient-derived mutations from the original cohort ^1^ *in vitro* (Figure 5A). Since E194, K276 and L278 are all located in α-helices of the 4HB domain (Figure 7A), their deletion would be predicted to induce a register shift in the alpha helix that is incompatible with folding of the 4HB domain. We confirmed that deletion of either E194, K276 or L278 leads to misfolding of MAST1^D^ ΔS90 and the formation of insoluble inclusion bodies in *E. coli* (Figure 7B). By contrast, the WT MAST1^D^ ΔS90 is almost entirely soluble (Figure 7B, S8A). Consistently, expression of a longer construct of MAST1 K276del (MAST1^DK^ ΔS90) in Sf9 insect cells resulted in aggregated protein with low yields (Figure 7C, S8B).

**Figure 7.**
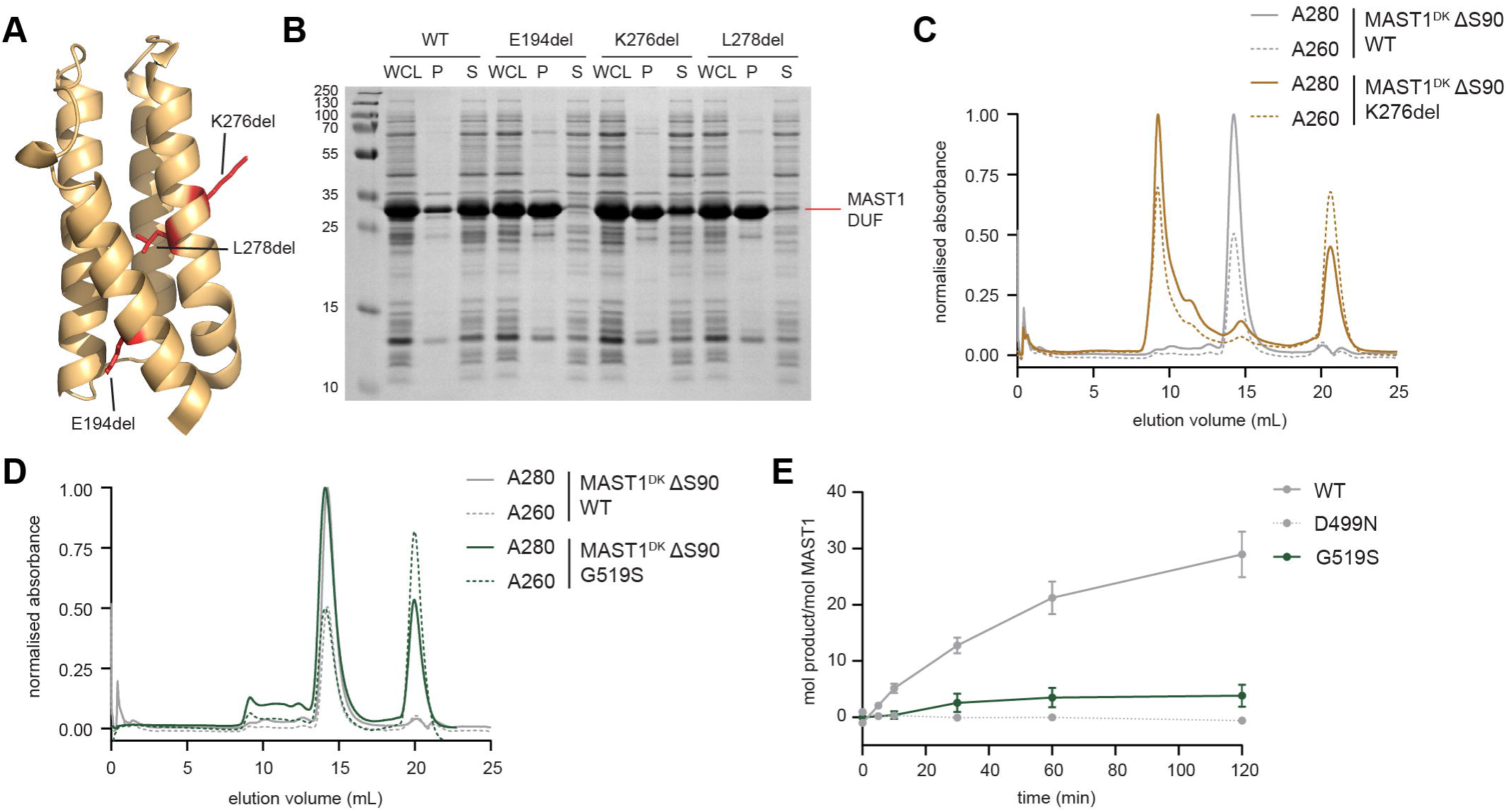
MCC-CH-CM mutations perturb MAST1 folding and kinase activity. A. Structure of MAST1 4HB determined by NMR (PDB ID 2M9X). Patient mutations residing in 4HB highlighted in red and labelled. B. Coomassie-stained SDS-PAGE gel of MAST1^D^ ΔS90 WT, E194del, K276del, or L278del, expressed in *E. coli*. WCL = whole cell lysate, P = pellet, S = supernatant. C. SEC elution profile of purified MAST1^DK^ ΔS90 WT and K276del from Sf9 insect cells, void volume = 9 mL. D. SEC elution profile of purified MAST1^DK^ ΔS90 WT and G519S from Sf9 insect cells. E. Radiometric kinase assay by MAST1^DK^ ΔS90 WT, kinase dead (D499N), or G519S on Tau. Data are presented as mean values ± S.E., *n* = 3.

Multiple patients have now been identified with a G517S mutation (mouse G519S), in the kinase domain. Although MAST1^DK^ ΔS90 bearing the G519S mutation was properly folded and monomeric by SEC (Figure 7D, S8C), it exhibited significantly lower kinase activity against Tau *in vitro*, albeit not completely inactive (Figure 7E). Taken together, these observations show that MCC-CH-CM mutations disrupt the folding and/or catalytic activity of MAST1.

## Discussion

We have discovered that MAST1 interacts with 14-3-3η in a phosphorylation-specific manner, mediated by S90 and S161. We also identified a group II PAK as the upstream kinase for MAST1 S90 phosphorylation, and S161 as a MAST1 autophosphorylation site. Finally, we provide evidence, both *in vitro* and *in vivo*, that the microtubule-associated protein Tau is a MAST1 substrate.

Our finding that PAK phosphorylates MAST1 sheds light on the broader biological pathways that MAST1 may be involved in. Like MAST1, PAK4 is highly expressed during embryogenesis, and is crucial in neurodevelopment, where it regulates the proliferation of neural progenitor cells ^42^, neuronal differentiation, and migration ^43^. This stems from PAK4’s role in the Cdc42-dependent reorganization of the actin cytoskeleton ^44–48^. Depletion of either Cdc42 or PAK4 results in cell polarity defects in a number of cell types ^49^, while knockout of PAK4 is embryonic lethal ^43^. PAK6 activity has been correlated with neurite outgrowth ^50^, and while PAK5/PAK6 double knockout mice are viable, they exhibit locomotor, learning and memory defects ^51^. Coordination of the actin cytoskeleton is crucial for axonal growth and steering. Actin and microtubule dynamics are coupled in the peripheral domain of the growth cone, together driving accurate axon navigation ^52,53^. Importantly, dysregulation of the growth cone can cause axonal rerouting, resulting in the disruption of axonal fiber tracts such as the corpus callosum ^54^. Here, Cdc42-activated group II PAKs may link actin polymerization and MAST1-mediated microtubule reorganization. However, the mechanisms that mediate the co-localization of MAST1 and PAK will require further investigation.

We have identified S214 of Tau as a MAST1 substrate. Tau binds to microtubules via its microtubule binding domain, encompassing the proline-rich domain (which contains S214) and the repeat domains, allowing Tau to rigidify and stabilize microtubules. Several studies have demonstrated the decreased microtubule binding capacity of Tau p-S214, reducing the stability of axonal microtubules ^36,55,56^. This effect is a combination of a decrease in microtubule binding affinity when Tau is phosphorylated ^55^, and the sequestration of Tau p-S214 by 14-3-3, which we and others ^40,41^ have observed. Interestingly, S214 has previously been identified as a cAMP-dependent protein kinase (PKA) phosphorylation site ^34,36,40,41,58^. Both PKA and the MAST kinases are AGC kinases, which recognize a conserved substrate consensus motif. It is therefore possible that MAST1 and PKA converge on the same substrate under different cellular conditions. It is also worth noting that, since MAST1 S161 and Tau S214 motifs are near-identical, it is perhaps not surprising that 14-3-3η is a common component in the signaling pathway. Curiously, hyperphosphorylation of S214 is also a marker of Alzheimer’s disease ^34,36^, though the significance of the connection to neurodevelopment is unclear. Our data also demonstrates that, at least *in vitro*, MAST1 is capable of phosphorylating Tau on S262, S324 and S356, all of which are reported phosphosites that inhibit microtubule binding ^37–39^. Nevertheless, these sites deviate from canonical AGC recognition motifs, so whether MAST1 is the physiologically relevant kinase *in vivo* will require further investigation. Whilst we have not yet explored the implications of MAST1-mediated Tau phosphorylation in the context of neurodevelopment, we hypothesize that MAST1 may negatively regulate microtubule polymerization in neuronal axons to ensure proper formation of the corpus callosum. Indeed, while the loss of Tau alone appears to have little impact on corpus callosum formation, a Tau and MAB1B double-knockout in mice causes a severe reduction in callosal size ^57^, beyond the more moderate agenesis phenotype observed in MAB1B-only knockout animals ^57,59^. This suggests a potential role of Tau in corpus callosum development, which may be partially redundant with MAP1B.

By combining these cellular insights with mechanistic insights into the molecular regulation of MAST1, we propose a model for MAST1 signaling (Figure 8). Activated Cdc42 recruits a group II PAK to the plasma membrane, whereupon it phosphorylates MAST1 on S90. MAST1 p-S90 specifically recruits 14-3-3η, expression of which is highly enriched in the brain and ubiquitously expressed in all brain structures ^60–63^, forming a complex with 1:2 stoichiometry. Since deletion of the S90 motif results in a loss of 14-3-3 binding to S161, we hypothesize that recruitment of 14-3-3η via p-S90 is required prior to S161 autophosphorylation and subsequent 14-3-3η binding. PAK-mediated phosphorylation of MAST1 S90 could therefore serve to pre-assemble a MAST1:14-3-3η complex. We hypothesize that the p-S90-bound complex is inactive, primed for activation by an unknown cofactor. The binding of a stimulatory cofactor to either MAST1 or the MAST1:14-3-3η complex would trigger MAST1 autophosphorylation at S161, thereby relieving autoinhibition. The consequences of p-S161 displacement from the kinase domain are two-fold: activation of the kinase, and sequestration of p-S161 by 14-3-3η, the latter of which may serve to both prevent reassociation of S161 and protect it from dephosphorylation. Activated MAST1 phosphorylates Tau on S214, causing the dissociation of Tau from microtubules. In short, Cdc42 signaling at the plasma membrane is transduced, through MAST1, to microtubule reorganization and changes in cell morphology.

**Figure 8.**
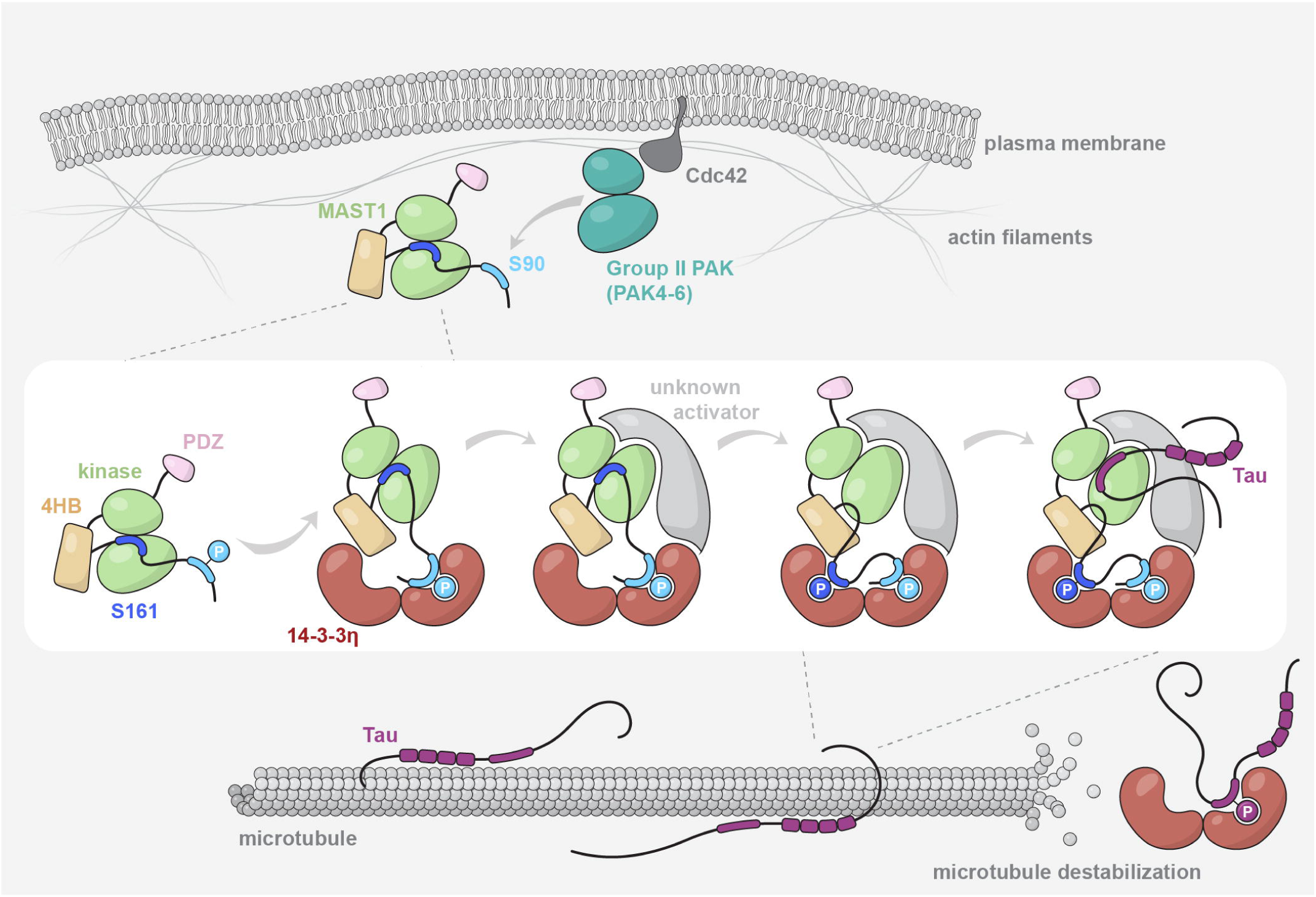
Model of MAST1 regulation. A group II PAK (PAK4-6) is activated by Cdc42 and phosphorylates MAST1 on S90. MAST1 p-S90 recruits 14-3-3η, forming an inactive trimeric complex, with S161 blocking MAST1’s substrate binding site. Upon binding of an unknown cofactor, the kinase domain of MAST1 adopts the active conformation and autophosphorylates S161, expelling it from the substrate binding site. p-S161 is sequestered by 14-3-3η, preventing dephosphorylation and reassembly with the kinase domain. Active MAST1 phosphorylates Tau on S214, resulting in its dissociation from microtubules, sequestration by 14-3-3η, and subsequent microtubule depolymerization.

While our findings imply a PAK-dependent coupling between actin and microtubule reorganization via MAST1, there are still many unanswered questions, particularly regarding the mechanism of activation of MAST1. Our data currently do not support a direct role for 14-3-3η in kinase activation, and MAST1 exhibits low activity *in vitro*. These observations beg the obvious question of precisely what role 14-3-3η plays in MAST1 regulation. Like MAST1, the oncoprotein BRAF possesses two regulatory phosphoserines, which simultaneously occupy the binding grooves of a 14-3-3 dimer. This trimeric complex maintains BRAF in an inactive conformation ^64,65^, which, upon RAS binding, is converted into an active complex with 2:2 stoichiometry ^64^. Whilst it is easy to envisage a similar mode of regulation for MAST1, both p-S90 and p-S161-mediated MAST1:14-3-3η complexes were observed to be trimeric, and a tetrameric complex has, so far, not been observed.

While the involvement of 14-3-3η in MAST1 signaling is clear, we speculate that there is another, as yet unidentified, factor that drives its activation. AGC kinases, such as MAST, are defined by a ∼50 amino acid C-terminal tail containing multiple regulatory motifs, including a hydrophobic motif that allosterically promotes the active conformation ^11^. In the closely related Dystrophia Myotonica Protein Kinase (DMPK) kinases, the hydrophobic motif is sandwiched in place by dimerization of the N-terminal capped helix bundle domain ^66–69^, while the Nuclear Dbf2-related (NDR) kinases are known to depend on MOB1 for activation via a similar mechanism ^70–73^. Currently, neither a MAST1-intrinsic domain nor a co-factor has been identified that could fulfil an analogous function. Although inspection of the MAST1 interactome in HEK293s (Figure 1C) failed to reveal any candidate activators by AlphaFold screening, identification of such a factor may depend on the correct cellular context, possibly in differentiated neurons.

Finally, this study provides important insights into the molecular consequences of MCC-CH-CM disease-associated mutations. We show that pathogenic mutations in MAST1 perturb folding and/or catalytic activity of the protein. Similar to the L278del mouse mutant, we report that the G519S mutation causes a notable decrease in the protein levels of MAST2, MAST3 and MAST4. Since MAST1 knock-out mice are phenotypically normal, this implies that MCC-CH-CM is the consequence of a dominant negative effect. As 14-3-3 proteins are obligate homodimers, the integration of mutant MAST1 has the potential to poison wild-type MAST complexes, though this would depend on hetero-tetramers of MAST and 14-3-3. Alternatively, the localization of mutant MAST1 to microtubules or titration of essential co-factors may inhibit signaling by wild-type MAST1.

In summary, MAST1 is a key regulator of neurodevelopment that potentially links group II PAKs, neuronal regulators of the actin cytoskeleton, to microtubule dynamics. However, several questions surrounding the precise mechanism of MAST1 activation, the origin of the dominant negative effect, and the stoichiometry of 14-3-3η binding remain. Answers to these questions are undoubtedly required to elucidate the physiological role of MAST1 in neurodevelopment as well as its pathological role in MCC-CH-CM.

## Acknowledgements

Proteomics analyses were performed on instruments of the Vienna BioCenter Core Facilities (VBCF). We acknowledge members of the Leonard lab for constructive feedback on the manuscript. This work was funded by Austrian Science Fund (FWF) grants P33066, P6212, P36724, and W1261 to T.A.L. We thank Boehringer Ingelheim for funding basic research and the core facilities at the Institute for Molecular Pathology, and Ludwig-Maximilians-University. P.H is supported by the Studienstiftung des deutschen Volkes.

## Author Contributions

S.A., P.H., D.A.K. and T.A.L. designed the experiments. S.A. performed all experiments reported in this manuscript with the exception of: proteomics analyses (W.C., D.A., M.H., K.M., G.D.); generation of mouse lines (P.H., R.T., M.F.M.R.); preparation of mouse brain samples for phosphoproteomics (P.H., R.T., M.F.M.R.); analysis of G519S mouse line (P.H.). D.A.K. and T.A.L. obtained funding for the project. S.A., P.H. D.A.K., and T.A.L. wrote the manuscript. All authors commented on the manuscript.

## Materials and Methods

### EGFP-MAST1 AP-MS

N-terminally EGFP-tagged MAST1^DKP^ constructs (WT, ΔS90, ΔS90 ΔS161, S90A, S161A, S90A S161A) were cloned into the pEGFP-C1 vector for expression in HEK293 cells and were transfected using Lipofectamine 3000 (Invitrogen). Media on the cells was exchanged 24 h post-transfection and cells were harvested by trypsinization 48 h post-transfection. Cells were lysed with three freeze/thaw cycles in liquid nitrogen, resuspended in lysis buffer (10 mM monosodium phosphate, 40 mM disodium phosphate, 10 mM NaCl, 50 mM NaF, 1 mM TCEP, 10 U Denarase, 1x protease inhibitor cocktail (Sigma P8849), 2 mM benzamidine, 2 mM MgCl2, 0.25% CHAPS, 10 mM betaglycerophosphate, 2 mM sodium pyrophosphate, 1 mM sodium orthovanadate, 1 mM EDTA), and incubated at 4°C for 30 min with rotation. Cleared lysates were obtained by centrifuging at 21,000 g, 4°C, for 15 min. Cleared lysates were incubated with GFP-Trap Magnetic beads (Chromotek), incubated at 4°C for 2 h while rotating, washed five times with wash buffer (10 mM monosodium phosphate, 40 mM disodium phosphate, 10 mM NaCl, 50 mM NaF, 1 mM TCEP), and the dry beads were snap-frozen in liquid nitrogen and stored at -80°C, for analysis by mass spectrometry.

### Fluorescence anisotropy

A 2x serial dilution series was made with purified protein (14-3-3η, 14-3-3θ, or MAST1^DK^ ΔS90 ΔS161) and mixed with 50 nM of N-terminally FITC-conjugated peptide and incubated at room temperature for 15 min. The following peptides were used in this study: DGRRWSLASLPSS (MAST1^85–97^), DGRRW{pS}LASLPSS (MAST1^85–97^ p-S90), RPRSRSLSPGRSP (MAST1^156–168^), RPRSR{pS}LSPGRSP (MAST1^156–168^ p-S161), PLGDP{pS}GRWARLS (MAST1 p-Scramble), RSRSL{pS}PGRSPSS (MAST1^158–170^ p-S163), PAVRPRSRSLSPGR (MAST1^153–166^ WT), PAVEPESRSLSPGR (MAST1^153–166^ R156E R158E), PAVEPESRSDSPGR (MAST1^153–166^ R156E R158E L162D), RSRTPSLPTPPTR (Tau^209–221^), and RSRTP{pS}LPTPPTR (Tau^209–221^ p-S214). Anisotropy was measured in a 384-well black polystyrene plate on the TECAN Spark Multi-Mode Microplate Reader.

### Radiometric kinase assay

For substrate phosphorylation assays, 100 nM of kinase was mixed with 25 μM of substrate in assay buffer (50 mM HEPES pH 7.4, 150 mM NaCl, 1 mM TCEP, 2 mM MgCl_2_, 2 mM ATP, 0.25% CHAPS, spiked with 1 μL [γ-32P] ATP/100 μL reaction) and incubated at 23°C for up to 2 h. Aliquots were taken from the reaction volume and quenched with 80 mM EDTA over a time course, then equal volumes of each sample were subjected to SDS-PAGE. The gels were washed three times in dH_2_0, dried, exposed to a phosphor screen for 4 h, and imaged with an Amersham Typhoon imager (Cytiva). Densitometry was performed using ImageJ to quantify the bands of phosphorylated substrate. An internal standard was made by spotting 0.5, 1, 5, and 20 pmol of ATP from the reaction onto a strip of Whatman paper. The standards were also measured with densitometry and used to calculate the total mol of phosphorylated substrate in each sample.

For autophosphorylation assays, 50 nM, 150 nM, and 500 nM of MAST1^DK^ ΔS90 (WT and D499N) was incubated in assay buffer (50 mM HEPES pH 7.4, 150 mM NaCl, 1 mM TCEP, 2 mM MgCl_2_, 2 mM ATP, 0.25% CHAPS, spiked with 1 μL [γ-32P] ATP/100 μL reaction) at 23°C for up to 2 h. Aliquots were taken from the reaction volume and quenched with 80 mM EDTA over a time course, then equal volumes of each sample were spotted onto a 0.45 μm nitrocellulose membrane. The membrane was washed five times with 20 mL of 75 mM H_3_PO_4_, and exposed and imaged as described above. An internal standard and densitometry were performed as described above.

### Analytical size exclusion chromatography

A Superdex 200 Increase 3.2/300 size exclusion column (Cytiva) was connected to an Äkta Pure with a Micro configuration (Cytiva) and equilibrated in buffer (50 mM HEPES pH 7.4, 150 mM NaCl, 1 mM TCEP) at 4°C. 50 μL of 30 μM protein samples were centrifuged at 21,000 g for 5 min prior to injection onto the column. For the MAST1^D^ p-S90 and 14-3-3η complex formed *in vitro*, equimolar concentrations of each protein were mixed and incubated for 30 min at room temperature prior to injection. 50 μL fractions were collected from each run for analysis by Coomassie-stained SDS-PAGE.

### Mass photometry

Purified protein samples were clarified after thawing by centrifugation at 21,000 g, 4°C for 5 min and diluted to 0.5 μM in freshly filtered buffer (50 mM HEPES pH 7.4, 150 mM NaCl, 1mM TCEP). Measurements of 60 s duration were recorded at a final concentration of 25-50 nM on microscopy coverslips that were cleaned by sonication in water and isopropanol. All measurements were recorded using a Two^MP^ mass photometer (Refeyn) and analyzed using Discover MP (Refeyn). Protein mass was calculated using a contrast-to-mass calibration with NativeMark Protein Standard (Invitrogen).

### Histological analysis of mice

Adult brains were isolated after transcardial perfusion with 0.9% NaCl followed by 4% PFA for 5 min at a flow rate of approx. 6 mL per minute. Brains were dissected and post-fixed in 4% PFA overnight at 4°C. Brains were washed with PBS and cryoprotected in 30% Sucrose in PBS. For sectioning, brains were embedded in Neg-50™ Medium (Richard-Allan Scientific) and sectioned into 40 µM-thick coronal brain sections using a sledge microtome. Nissl stainings were carried out via incubation in a 0.25% solution of Cresyl Violet acetate (Sigma, #C5042), followed by dehydration in a graded ethanol series (30%, 70%, 95%, 100%; 3 min, 2 min, 30 s, 10 s) and xylol (2 x 2 min) and mounted using DPX mountant (Fluka, #44581). Slides were left to solidify overnight at RT and images were acquired using a Mirax slide scanner (Zeiss). The thickness of the corpus callosum was measured at Bregma + 0.86 mm by performing 3 parallel measurements near the midline position using ImageJ. The cortical thickness was measured at Bregma -1.82 mm through bilateral measurements of the somatosensory cortex at 3 matched positions. Averages were taken and used for analysis.

## Notes

https://www.ebi.ac.uk/pride/

### Competing Interest Statement

The authors have declared no competing interest.

