## Supplementary Information for "A complex of MAST1 and 14-3-3η regulates Tau phosphorylation in the developing cortex"

Contains: Supplementary Figures 1-8

Supplementary Table 1

Supplementary Methods

Supplementary References

#### Supplementary Figures

##### Supplementary Figure 1. MAST1 interacts with 14-3-3 proteins via S90 and S161.

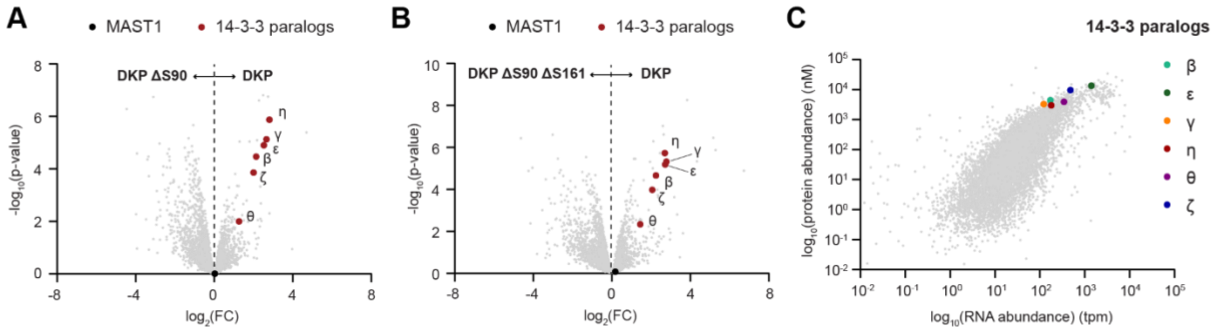


A. IP-MS for EGFP-MAST1$^{DKP}$ (right) vs EGFP-MAST1$^{DKP}$ ΔS90 (left), *n* = 3.

B. IP-MS for EGFP-MAST1$^{DKP}$ (right) vs EGFP-MAST1$^{DKP}$ ΔS90 ΔS161 (left), *n* = 3.

C. Scatter plot showing RNA and protein abundance of all six 14-3-3 paralogs identified in Figure 1C (highlighted by large colored dots) overlayed on all detected RNA/protein, in HEK293 cells [12].

Supplementary Figure 2. Mass spectra of all protein constructs employed in this study.

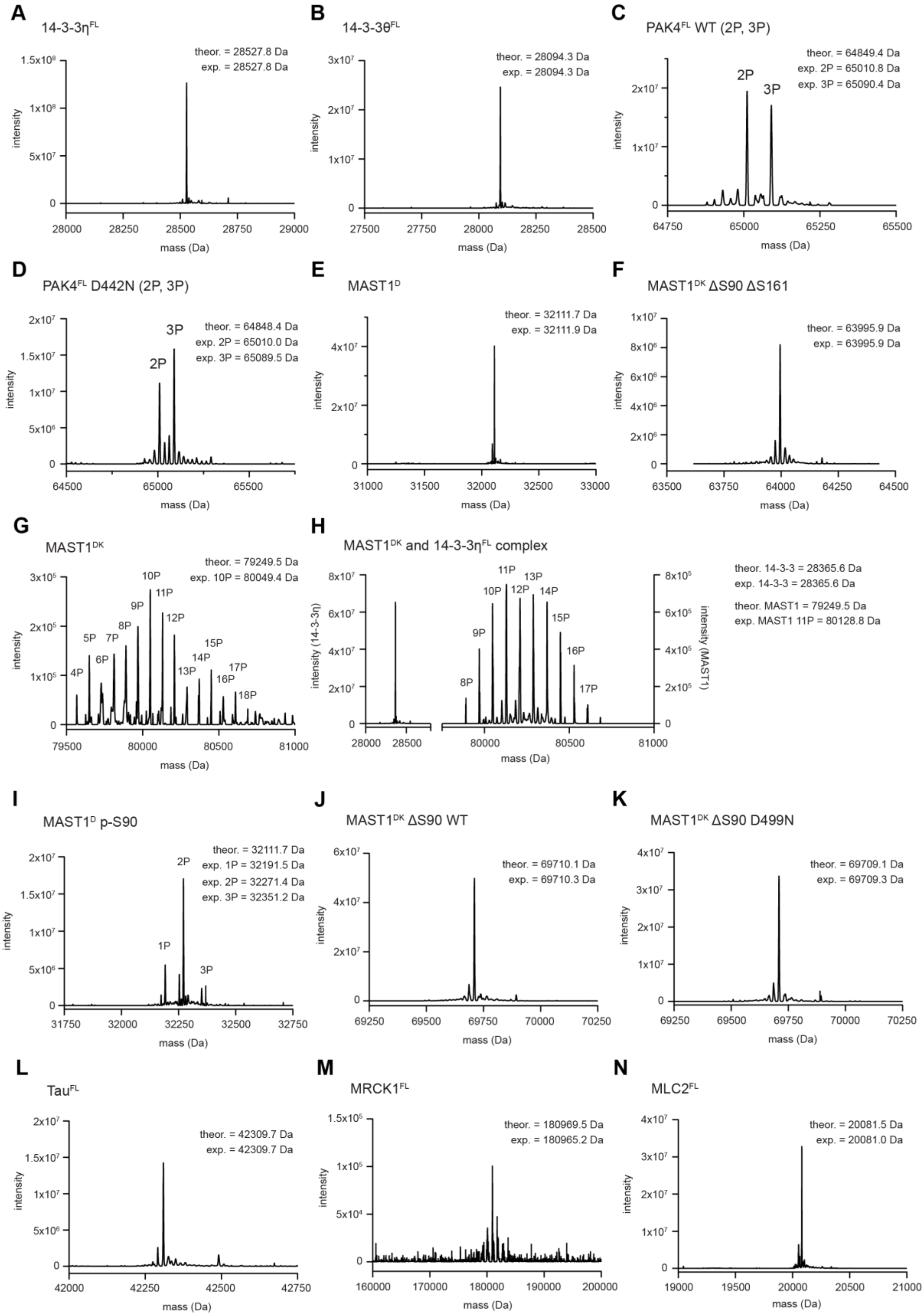

**O**

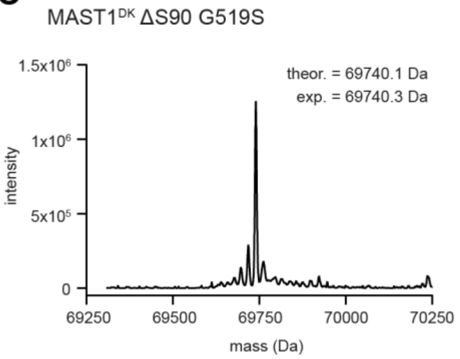


A. Mass spectrum of purified 14-3-3η$^{FL}$.

B. Mass spectrum of purified 14-3-3θ$^{FL}$.

C. Mass spectrum of purified PAK4$^{FL}$ WT.

D. Mass spectrum of purified PAK4$^{FL}$ D442N.

E. Mass spectrum of purified MAST1$^{D}$.

F. Mass spectrum of purified and dephosphorylated MAST1$^{DK}$ ΔS90 ΔS161.

G. Mass spectrum of purified MAST1$^{DK}$.

H. Mass spectrum of co-purified MAST1$^{DK}$ and 14-3-3η$^{FL}$ complex.

I. Mass spectrum of purified MAST1$^{D}$ after phosphorylation by PAK4$^{FL}$ WT.

J. Mass spectrum of purified and dephosphorylated MAST1$^{DK}$ ΔS90 WT.

K. Mass spectrum of purified and dephosphorylated MAST1$^{DK}$ ΔS90 D499N.

L. Mass spectrum of purified Tau$^{FL}$.

M. Mass spectrum of purified MRCK1$^{FL}$.

N. Mass spectrum of purified MLC2$^{FL}$.

O. Mass spectrum of purified and dephosphorylated MAST1$^{DK}$ ΔS90 G519S.

Supplementary Figure 3. MAST1 interacts with 14-3-3 proteins via S90 and S161.

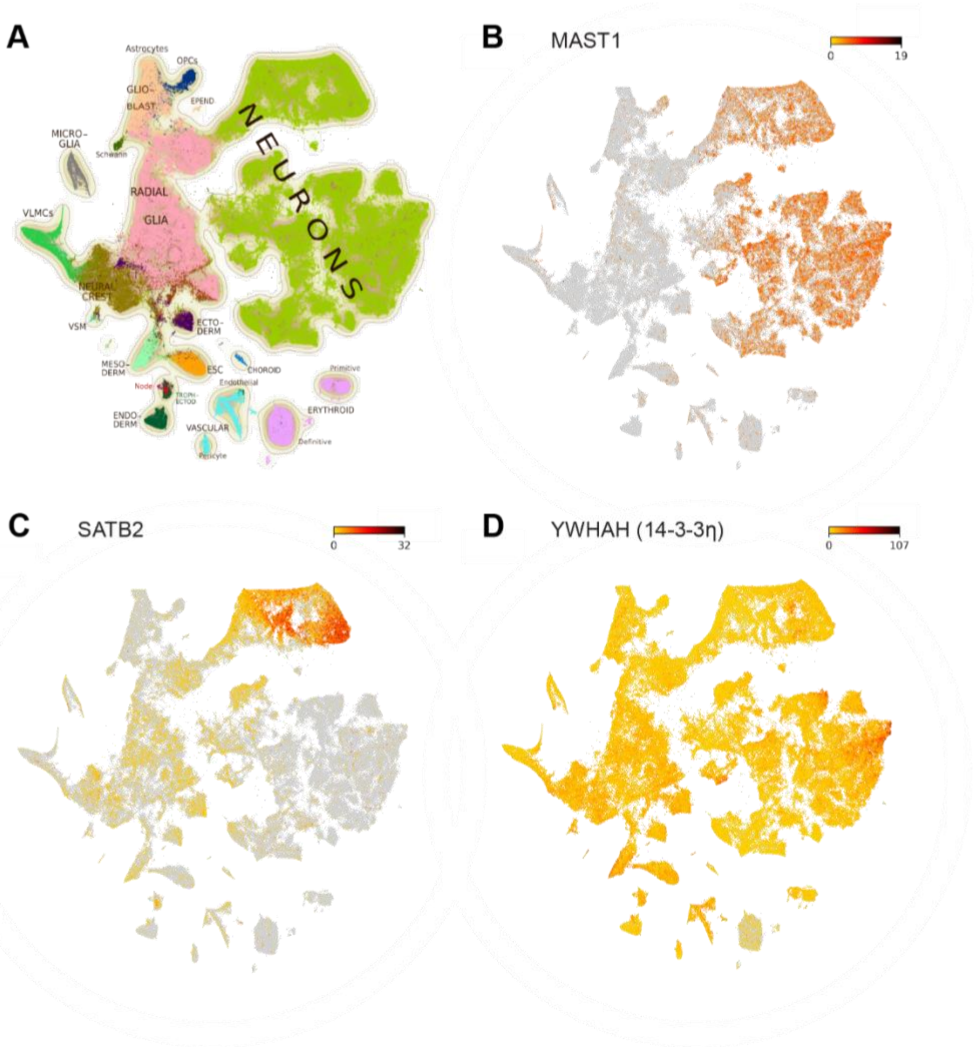


A. t-SNE plot of single-cell RNA sequencing data from embryonic mouse brains between embryonic stage E7-E18, colored by major cell types [14].

B. MAST1 expression levels overlayed on t-SNE plot from panel A.

C. SATB2 expression levels overlayed on t-SNE plot from panel A.

D. YWHAH (14-3-3η) expression levels overlayed on t-SNE plot from panel A.

#### Supplementary Figure 4. PAK4 phosphorylates MAST1 on S90.

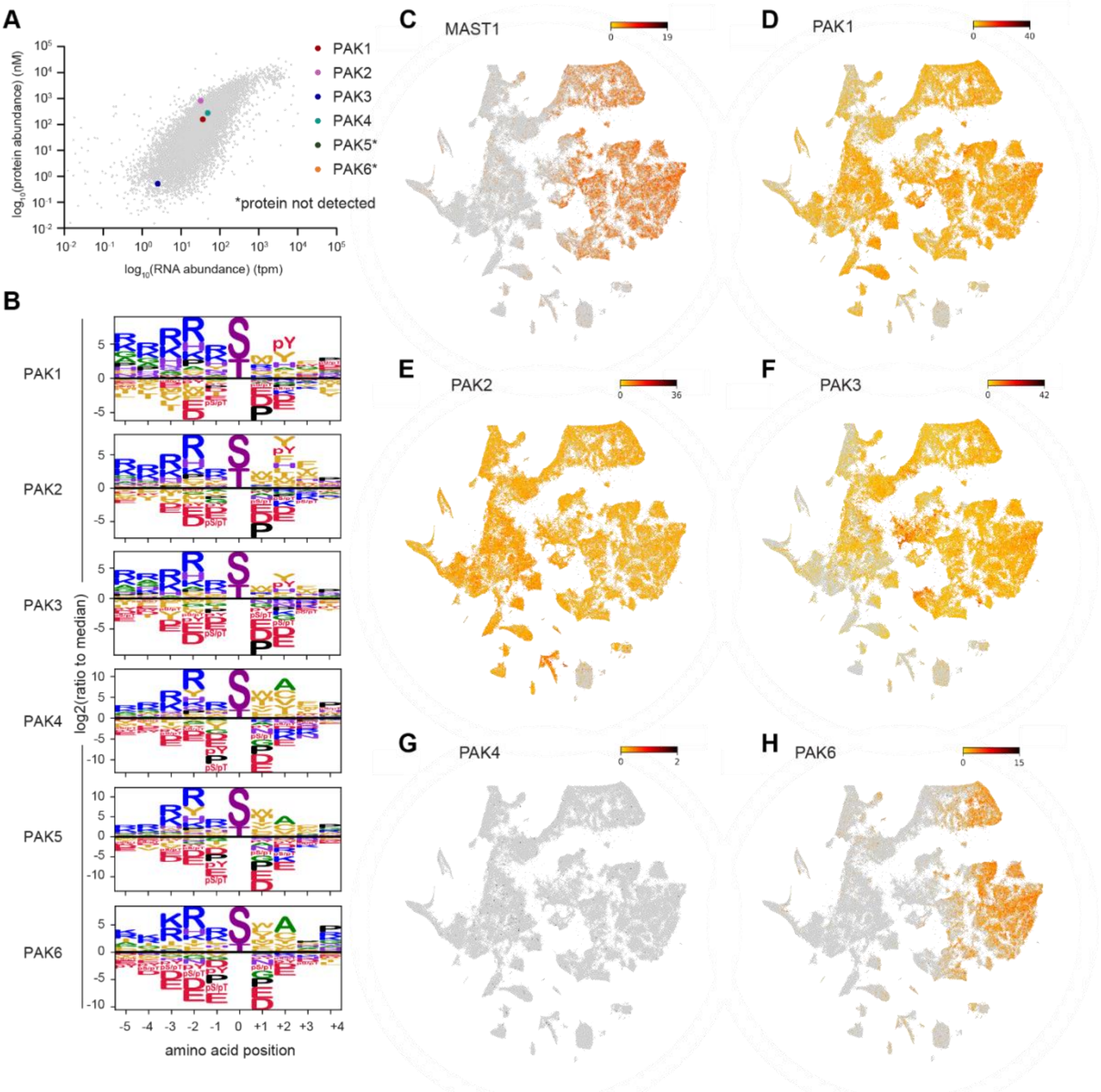


A. Scatter plot showing RNA and protein abundance of all PAKs (highlighted by large colored dots) overlayed on all detected RNA/protein, in HEK293 cells [12]. Both PAK5 and PAK6 protein was not detected, and are not plotted.

B. Substrate motif sequence logos of PAK1-6 from P-5 to P+4, where position 0 is the phosphorylatable residue [15].

C. MAST1 expression levels overlayed on t-SNE plot from Figure S3A.

D. PAK1 expression levels overlayed on t-SNE plot from Figure S3A.

E. PAK2 expression levels overlayed on t-SNE plot from Figure S3A.

F. PAK3 expression levels overlayed on t-SNE plot from Figure S3A.

G. PAK4 expression levels overlayed on t-SNE plot from Figure S3A.

H. PAK6 expression levels overlayed on t-SNE plot from Figure S3A.

Supplementary Figure 5. MAST1 S161 is an autoregulatory motif.

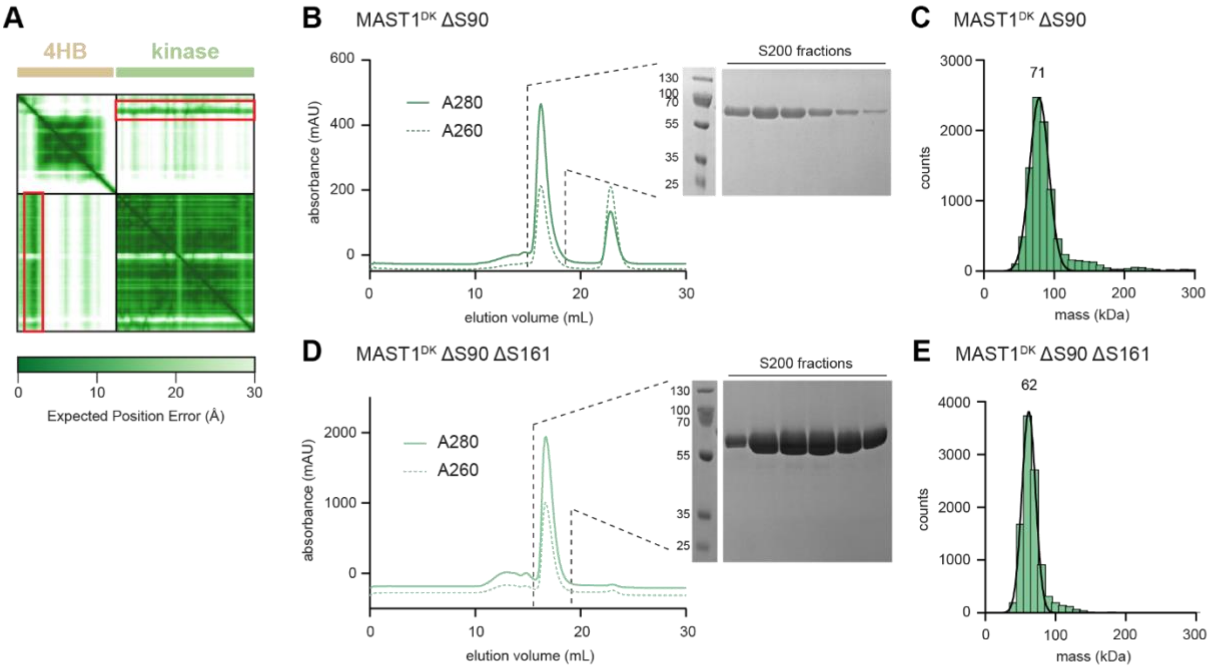


A. Pair alignment error (PAE) plot of MAST1 kinase domain and DUF. Region spanning the S161 sequence motif outlined in red box.

B. Size exclusion chromatography (SEC) elution profile of purified MAST1$^{DK}$ ΔS90. Coomassie-stained SDS-PAGE gel of SEC fractions spanning elution peak shown in inset.

C. Mass photometry of purified MAST1$^{DK}$ ΔS90. Theoretical mass = 70 kDa, observed mass = 71 kDa.

D. SEC elution profile of purified MAST1$^{DK}$ ΔS90 ΔS161. Coomassie-stained SDS-PAGE gel of SEC fractions spanning elution peak shown in inset.

E. Mass photometry of purified MAST1$^{DK}$ ΔS90 ΔS161. Theoretical mass = 64 kDa, observed mass = 62 kDa.

Supplementary Figure 6. MAST1 p-S90 or p-S161 forms a complex with 14-3-3η *in vitro*.

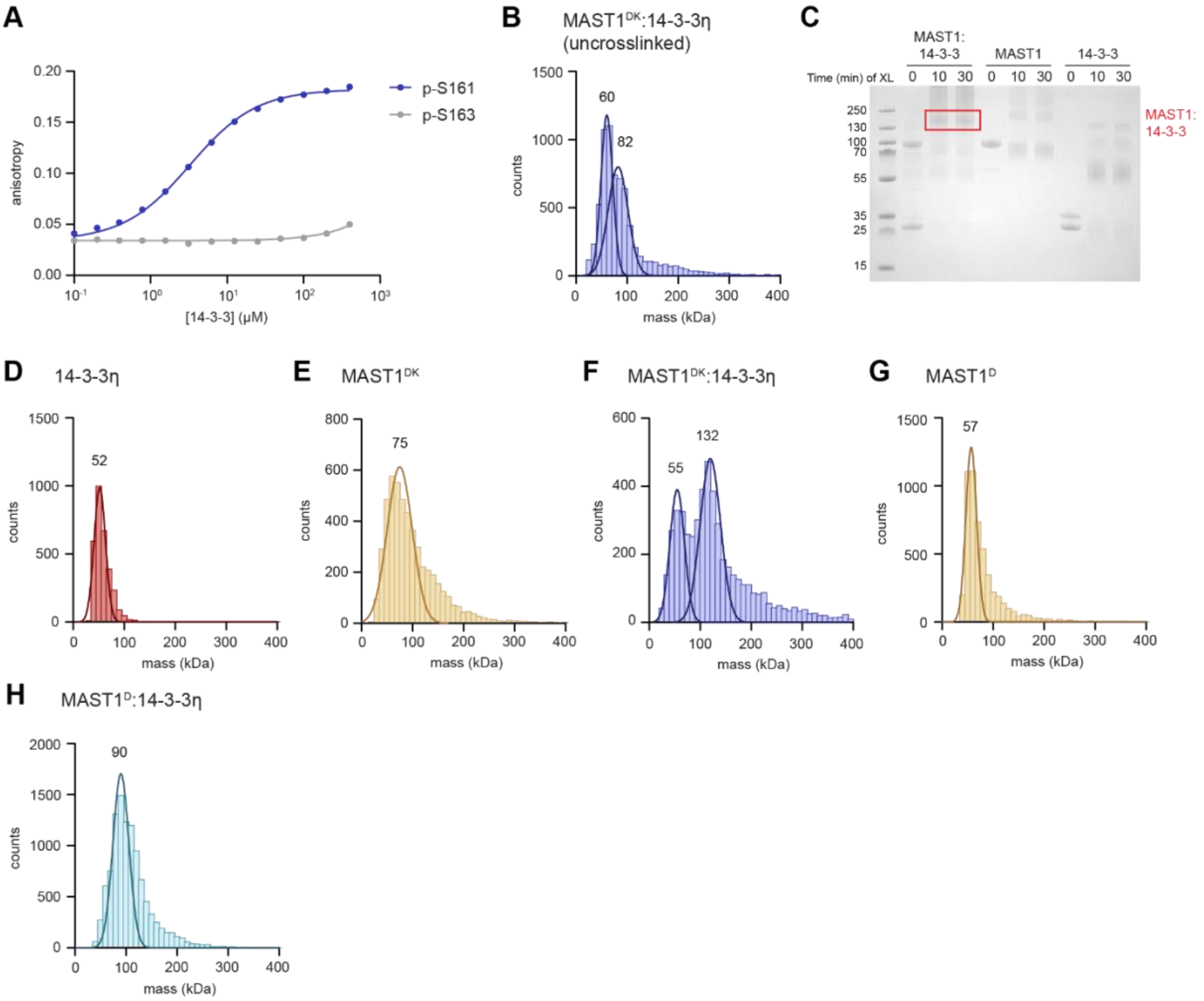


A. Binding curves for 14-3-3η and MAST1 peptides spanning S161 or S163, measured by fluorescence anisotropy. Data are presented as mean values ± S.E., *n* = 3. Data were fit with a one-site binding model and the $K_d$ derived from curve fitting.

B. Mass photometry of co-purified $MAST1^{DK}$ and 14-3-3η complex.

C. Coomassie-stained SDS-PAGE gel of $MAST1^{DK}$ and 14-3-3η complex, $MAST1^{DK}$ and 14-3-3η after 0, 10, and 30 min of glutaraldehyde crosslinking. Crosslinked trimeric complex outlined in red box.

D. Mass photometry of purified 14-3-3η after 10 min of glutaraldehyde crosslinking.

E. Mass photometry of purified MAST1$^{DK}$ after 10 min of glutaraldehyde crosslinking.

F. Mass photometry of co-purified MAST1$^{DK}$ and 14-3-3η complex after 10 min of glutaraldehyde crosslinking.

G. Mass photometry of purified MAST1$^{D}$ p-S90.

H. Mass photometry of *in vitro*-formed MAST1$^{D}$ p-S90 and 14-3-3η complex after 10 min of glutaraldehyde crosslinking.

#### Supplementary Figure 7. Tau is a MAST1 substrate.

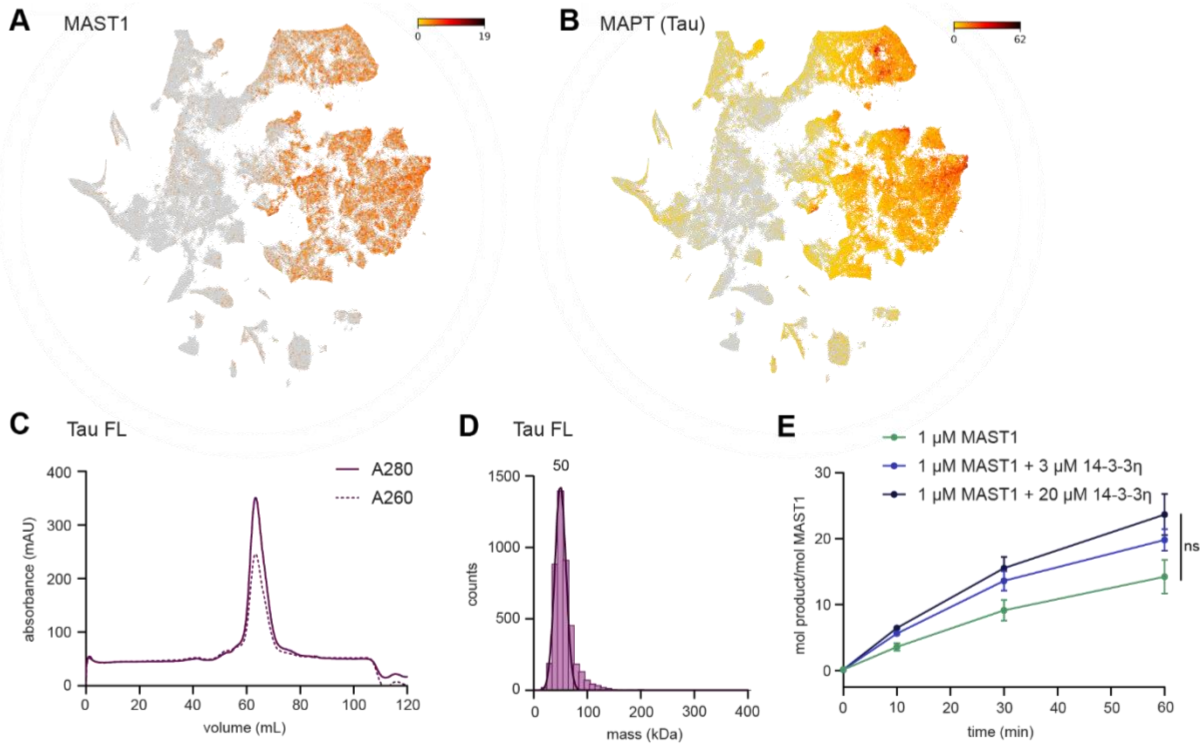


A. MAST1 expression levels overlayed on t-SNE plot from Figure S3A.

B. MAPT (Tau) expression levels overlayed on t-SNE plot from Figure S3A.

C. SEC elution profile of purified Tau.

D. Mass photometry of purified Tau. Theoretical mass = 42 kDa, observed mass = 50 kDa.

E. Radiometric kinase assay by $MAST1^{DK}$, $MAST1^{DK}$ and 14-3-3η complex, or $MAST1^{DK}$ and 14-3-3η complex supplemented with excess of purified 14-3-3η, on Tau ($n$ = 3, p = 0.0642, two-way repeated-measures ANOVA). Data are presented as mean values ± S.E.

#### Supplementary Figure 8. MCC-CH-CM mutations perturb MAST1 folding and kinase activity.

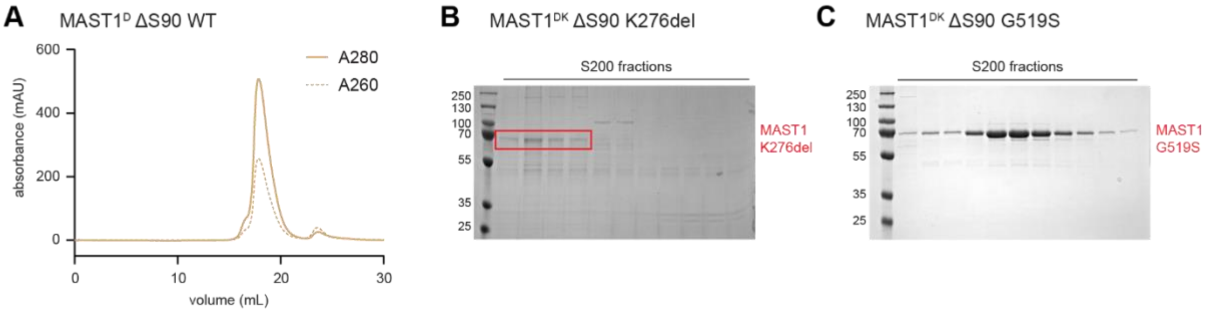


A. SEC elution profile of purified MAST1$^{D}$ $\Delta$S90 WT.

B. Coomassie-stained SDS-PAGE gel of SEC fractions spanning elution peak of MAST1$^{DK}$ $\Delta$S90 K276del. Bands representing MAST1 outlined in red box.

C. Coomassie-stained SDS-PAGE gel of SEC fractions spanning elution peak of MAST1$^{DK}$ $\Delta$S90 G519S.

### Supplementary Tables

#### Supplementary Table 1. Oligonucleotide sequences for generation of G519S mouse model

| | |
|---|---|
| mMAST1-G517S sgRNA target sequence | CACATCAAGCTCACGGATTTTGG |
| mMAST1-G517S ssDNA repair template | CCTGAGATCCCCACCTGTGCCTACAGCCTCCTTATCACCTCCATGGGTCACA TCAAGCTCACAGATTTCGGCCTCTCCAAGATGGGGCTCATGAGCCTCACCAC CAACTTATATGAAGGCCACATCGAGAAGGACGCCCC |
| mMAST1-G517S genotyping primer fw | CTCAAGCCAGACAAGTGAG |
| mMAST1-G517S genotyping primer rv | CCTGTTTGTCCAGGAACTC |

### Supplementary Methods

#### Protein expression and purification

##### *Sf9 expression*

All proteins expressed are derived from *Mus musculus* sequence unless otherwise stated. TEV-cleavable N-terminal 10xHis-tagged constructs of $MAST1^{DK}$, $MAST1^{DK}$ ΔS90, $MAST1^{DK}$ ΔS90 ΔS161, and $PAK4^{FL}$, 3C-cleavable N-terminal GST-tagged constructs of 14-3-3η, and a TEV-cleavable N-terminal StrepII-tagged construct of *Homo sapeins* $MRCK\alpha^{FL}$ were cloned into the pFastBacDual vector and expressed in baculovirus-infected *Spodoptera frugiperda* (Sf9) insect cells. Mutant constructs were cloned using site-directed mutagenesis. Sf9 cell pellets from 1 L of expression culture were lysed in 100 mL lysis buffer (50 mM HEPES pH 7.4, 150 mM NaCl, 1 mM TCEP, 2 mM benzamidine, 2 mM $MgCl_2$, 1x protease inhibitor cocktail (Sigma P8849), 1 kU Denarase (c-LEcta)). 20 mM imidazole was added for constructs purified by Ni-NTA affinity chromatography, 0.25% CHAPS was added for constructs purified by GST-beads, and phosphatase inhibitors (50 mM NaF, 10 mM betaglycerophosphate, 2 mM sodium pyrophosphate, 1 mM sodium orthovanadate) was added for $MAST1^{DK}$/14-3-3η co-purification to retain phosphates. Cleared lysates were obtained by centrifugation (38,000 g, 4°C, 30 min).

##### *BL21 expression*

All proteins expressed are derived from *Mus musculus* sequence unless otherwise stated. TEV-cleavable N-terminal 6xHis-tagged constructs of $MAST1^{D}$, 14-3-3η, and 14-3-3θ were cloned into the pHisII vector, an N-terminal 6xHis-tagged construct of $Tau^{FL}$ (isoform F) was cloned into the pET28b vector, and a TEV-cleavable N-terminal GST-tagged construct of *Homo sapiens* $MLC2^{FL}$ was cloned into the pGST2 vector, for expression in BL21 Star *E. coli*. The cells were grown at 37°C to an OD of 0.7 before induction with 250 µM IPTG and overnight expression at 18°C. BL21 cell pellets from 1 L of expression culture were lysed in 50 mL lysis buffer (50 mM HEPES pH 7.4, 150 mM NaCl, 1 mM TCEP, 2 mM benzamidine, 2 mM $MgCl_2$, 1x protease inhibitor cocktail (Sigma P8849), 0.5 kU Denarase (c-LEcta), 5 mg lysozyme). 20 mM imidazole and 0.25% CHAPS was added as described above. Lysates were sonicated three times for 3 min at 50% amplitude and cleared lysates were obtained by centrifugation (38,000 g, 4°C, 20 min).

##### *Purification*

For Ni-NTA affinity purification, cleared lysates were run over a HisTrap FF column (Cytiva), washed with buffer A (50 mM HEPES pH 7.4, 150 mM NaCl, 1 mM TCEP) followed by buffer A with 2% buffer B (buffer A + 1 M imidazole), and eluted with buffer A with 30% buffer B. The eluate was cleaved and dialyzed overnight at 4°C with 8.8 nmol TEV protease (purified in house) in 50 mM HEPES pH 7.4, 50 mM NaCl, and 1 mM TCEP. Constructs requiring dephosphorylation were treated with 2.3 nmol of lambda-phosphatase (purified in house) and 0.1 mM $MnCl_2$.

For GST affinity purification, cleared lysates were incubated with 3 mL glutathione sepharose beads (Cytiva) for 3 hours at 4°C. The beads were washed three times with buffer A (50 mM HEPES pH 7.4, 150 mM NaCl, 1 mM TCEP, 0.25% CHAPS) and the construct was cleaved off the beads overnight at 4°C with either 8.8 nmol TEV protease (purified in house) or 11.1 nmol 3C protease (purified in house).

For StrepII affinity purification, cleared lysates were run over a StrepTrap HP column (Cytiva), washed with buffer A (50 mM HEPES pH 7.4, 150 mM NaCl, 1 mM TCEP), and eluted with buffer B (buffer A + 2.5 mM d-desthiobiotin). The eluate was cleaved overnight at 4°C with 8.8 nmol TEV protease (purified in house).

Following affinity purification, constructs underwent ion-exchange purification. Proteins that were not dialyzed to 50 mM NaCl were first diluted to make a final concentration of 50 mM NaCl, then run over either a HiTrap Q HP or HiTrap SP HP column (Cytiva), washed with buffer A (50 mM HEPES pH 7.4, 1 mM TCEP), and eluted with a gradient of buffer B (50 mM HEPES pH 7.4, 1 M NaCl, 1 mM TCEP). The peak fractions were pooled, concentrated, and injected onto a size exclusion column (Cytiva) equilibrated in 50 mM HEPES pH 7.4, 150 mM NaCl, 1 mM TCEP. S200 and S75 columns were used for constructs >30 kDa and <30 kDa respectively, and 10/300 and 16/600 columns were used for yields >5 mg and <5 mg respectively. For $MAST1^{D}$ requiring phosphorylation on S90, it was incubated with purified $PAK4^{FL}$ at room temperature for 2 hours prior to injection on the size exclusion column. The peak fractions were pooled, concentrated, and stored at -80°C.

#### Cell culture

HEK293 cells were maintained in DMEM supplemented with 10% FBS. Cells were tested negative for *Mycoplasma* contamination using Eurofins Genomics Mycoplasmacheck service.

#### Intact mass spectrometry

Protein sample was diluted in dH2O to 4 ng/µL and dithiothreitol was added to a final concentration of 25 mM. 8 ng (2 µL) of protein was loaded on an XBridge Protein BEH C4 column (2.5 µm particle size, dimensions 2.1 mm x 150 mm; Waters) using a Vanquish™ Horizon UHPLC system (Thermo Scientific) with a working temperature of 50 °C, 0.1% formic acid (FA) as solvent A, 100% acetonitrile, 0.08% FA as solvent B. Proteins were separated on a 6 min step gradient from 12 to 72% solvent B at a flow rate of 250 µL/min and analyzed on a Synapt G2-Si coupled via a ZSpray ESI source (Waters). Data were recorded with MassLynx V 4.2 (Waters) and analyzed using the MaxEnt 1 process to reconstruct the uncharged average protein mass.

#### Tandem mass spectrometry

##### *Sample preparation*

For MAST1$^{D}$, MAST1$^{DK}$ ΔS90, and MAST1$^{DK}$:14-3-3η, ~5 µg protein in 1.5 µL was diluted with 8.5 µL of 8 M urea, 50 mM ammonium bicarbonate (ABC). For Tau$^{FL}$, ~5 µg of protein in 2.5 µL was mixed with 10% sodium deoxycholate (SDC) and 50 mM ABC to reach 2% SDC. Proteins were reduced with 0.4 µL of 250 mM dithiothreitol (DTT) for 30 min at room temperature (50°C for Tau$^{FL}$) and alkylated by adding 0.4 µL of 500 mM IAA and incubating for 30 min at room temperature in the dark. The remaining iodoacetamide (IAA) was quenched by adding 0.2 µL of 250 mM DTT for 10 min. For MAST1$^{D}$, MAST1$^{DK}$ ΔS90, and MAST1$^{DK}$:14-3-3η, the sample was diluted to below 1 M urea with 70 µL of 50 mM ABC. For Tau$^{FL}$, the sample was diluted to 1% SDC with 10 µL of 100 mM ABC. Proteins were digested by adding ~160 ng trypsin (Trypsin Gold, Promega) and incubated at 37°C overnight. The digest was stopped by the addition of 10% trifluoroacetic acid to a final concentration of 0.5% (1% for Tau$^{FL}$). For Tau$^{FL}$, samples were centrifuged at 18,500 x g for 10 min to remove the SDC, the supernatant was transferred to a new tube, 10% TFA was added to reach a final concentration of 2%, and the sample was centrifuged again at 18,500 x g for 10 min. The resulting supernatant was desalted using C18 Stagetips [1].

For EGFP-MAST1 AP-MS bead samples, the beads were resuspended in 30 µL of 2 M urea and 50 mM ammonium bicarbonate. Disulfide bonds were reduced with 1.2 µL of 250 mM DTT for 30 min at room temperature before adding 1.2 µL of 500 mM IAA and incubating for 30 min at room temperature in the dark. The remaining iodoacetamide was quenched with 0.6 µL of 250 mM DTT for 10 min. Proteins were digested with 150 ng LysC (mass spectrometry grade, FUJIFILM Wako chemicals) and trypsin (Trypsin Gold, Promega) mix at room temperature for 90 min. The supernatant was transferred to a new tube and the beads were rinsed with 30 µL of 2 M urea and 50 mM ammonium bicarbonate. 60 µL of 50 mM ammonium bicarbonate was added to the tube, before digestion with 150 ng LysC/trypsin mix at 37°C for 5 h. The digest was stopped by the addition of TFA to a final concentration of 0.5%, and the peptides were desalted using C18 Stagetips [1].

*Liquid chromatography-mass spectrometry analysis*

For MAST1$^{D}$, MAST1$^{DK}$ ΔS90, MAST1$^{DK}$:14-3-3η, and Tau$^{FL}$, LC-MS analysis was performed on a Vanquish Neo UHPLC system (Thermo Scientific) coupled to an Orbitrap Exploris 480 mass spectrometer (Thermo Scientific). The system was equipped with a Nanospray Flex ion source (Thermo Scientific), coated emitter tips (PepSep, MSWil), and a Butterfly Portfolio Heater (Phoenix S&T). Peptides were loaded onto a trap column (Acclaim PepMap 100 C18 HPLC Column, 20 mm × 0.1 mm, 5 µm particle size, Thermo Scientific) using 0.1% TFA as mobile phase, and separated on an analytical column (Acclaim PepMap 100 C18 HPLC Column, 50 cm × 75 µm, 2 µm particle size, Thermo Scientific), applying a linear gradient starting with a mobile phase of 98% solvent A (0.1% FA) and 2% solvent B (80% acetonitrile, 0.08% FA), increasing to 35% solvent B over 60 min at a flow rate of 230 nL/min. The analytical column was heated to 30°C.

The mass spectrometer was operated in data-dependent acquisition (DDA) mode, with 2 s MS1 cycle time. For MAST1$^{D}$, MAST1$^{DK}$ ΔS90, and MAST1$^{DK}$:14-3-3η, survey scans were acquired from 375-2000 m/z with lock mass enabled, normalized AGC target of 300%, resolution of 120,000. For Tau$^{FL}$, survey scans were acquired from 340-1500 m/z with lock mass enabled, normalized AGC target of 300%, resolution of 60,000. The most intense precursor ions (charge states +2 to +6) were selected for fragmentation using an isolation window of 1.2-1.4 m/z. Selected ions were analyzed with a maximum fill time of 100 ms, normalized AGC target of 200%, and resolution of 30,000 after HCD fragmentation with normalized collision energy of 28-30%. Monoisotopic precursor selection (MIPS) was set to “peptide” mode, the intensity threshold to 2.5 x $10^4$ (1 x $10^4$ for $Tau^{FL}$), and selected precursors were dynamically excluded for 20 seconds with isotope exclusion enabled.

For EGFP-MAST1 AP-MS samples, LC-MS analysis was performed on an UltiMate 3000 RSLCnano LC system (Thermo Scientific) coupled to a timsTOF HT (Bruker). The system was equipped with a CaptiveSpray ion source (Bruker), and a Butterfly Portfolio Heater (Phoenix S&T). Peptides were loaded onto a trap column (Acclaim PepMap 100 C18 HPLC Column, 20 mm × 0.1 mm, 5 µm particle size, Thermo Scientific) using 0.1% TFA as mobile phase, and separated on an analytical column (Aurora Ultimate XT C18, 25 cm × 75 µm, 1.7 µm particle size, IonOpticks), applying a linear gradient starting with a mobile phase of 98% solvent A (0.1% FA) and 2% solvent B (80% acetonitrile, 0.08% FA), increasing to 35% solvent B over 60 min at a flow rate of 300 nL/min. The analytical column was heated to 50°C.

The mass spectrometer was operated in data-independent acquisition (DIA) parallel accumulation serial fragmentation (PASEF) mode. MS2 data were acquired with eight PASEF scans per duty cycle, each containing three m/z windows. The m/z window widths were adjusted based on the expected precursor density, covering a total range of 300–1200 m/z. The ion mobility range was set to 0.64-1.42 V*s/cm, and the accumulation and ramp time was set to 100 ms. TIMS elution voltages were calibrated linearly to obtain the reduced ion mobility coefficients (1/K0) using three Agilent ESI-L Tuning Mix ions (m/z 622, 922 and 1,222). Collision energy for fragmentation was scaled linearly with precursor mobility (1/K0), ranging from 20 eV (at 1/K0 = 0.6 V*s/cm) to 59 eV at (at 1/K0 = 1.6 V*s/cm).

##### *Mass spectrometry data analysis*

For $MAST1^{D}$, $MAST1^{DK}$ ΔS90, $MAST1^{DK}$:14-3-3η, and $Tau^{FL}$, MS raw data were analyzed with FragPipe (22.0), using MSFragger (4.1) [2], IonQuant (1.10.27) [3], and Philosopher (5.1.1) [4]. The default FragPipe workflow for label free quantification (LFQ-MBR) was used, except “Normalize intensity across runs” was turned off. Cleavage specificity was set to Trypsin/P, with two missed cleavages allowed. The protein FDR was set to 1%. Carbamidomethyl was used as fixed cysteine modification; methionine oxidation, phosphorylation at STY and protein N-terminal acetylation were specified as variable modifications. PTMProphet was used for the PTM site localization. MS2 spectra were searched against the *Spodoptera frugiperda* protein sequences from Uniprot (Taxon ID: 7108, release 2024_01), concatenated with a database of 379 common laboratory contaminants (release 2023.03, https://github.com/maxperutzlabs-ms/perutz-ms-contaminants), and protein sequences for the constructs.

For EGFP-MAST1 AP-MS samples, MS raw data were processed with Spectronaut (19.0 or 19.2, Biognosys). The library-free DirectDIA+ workflow was employed for analysis of the raw files, the *H. sapiens* 1 protein per gene reference proteome from Uniprot (Proteome ID: UP000005640, release 2024_01), concatenated with a database of 379 common laboratory contaminants (release 2023.03, https://github.com/maxperutzlabs-ms/perutz-ms-contaminants), and sequences for the constructs. The cleavage specificity was set to full trypsin specificity (Trypsin/P), with two missed cleavages allowed. Carbamidomethyl was used as fixed cysteine modification; methionine oxidation and protein N-terminal acetylation were specified as variable modifications. Cross-run normalization was disabled, and all other settings were used at their default values.

##### *Post-processing with in-house pipeline*

Computational analysis was performed using Python and the in-house developed Python library MsReport (0.0.26) [5]. Only non-contaminant proteins identified with a minimum of two peptides and being quantified once were considered for further analysis. LFQ protein intensities were log2-transformed and normalized across samples using the ModeNormalizer from MsReport. The ModeNormalizer method involves calculating log2 protein ratios for all pairs of samples and determining normalization factors based on the modes of all ratio distributions. Missing values were imputed by drawing random values from a normal distribution. Sigma and mu of this distribution were calculated per sample from the standard deviation and median of the observed log2 protein intensities ($\mu$ = median sample LFQ intensity – 1.8 standard deviations of the sample LFQ intensities, $\sigma = 0.3 \times$ standard deviation of the sample LFQ intensities). iBAQ intensities were calculated by dividing protein intensities by the number of theoretically observable tryptic peptides between 6 and 30 amino acids. To estimate the relative protein abundances within each sample, the iBAQ intensities were normalized by dividing them by the total sum of iBAQ intensities for all proteins in the sample. For the EGFP-MAST1 AP-MS samples, statistical analysis was performed using the linear models for microarray analysis (limma) v.3.54.2 [6] package in R. Moderated t-statistics were calculated using the limma-trend method, and multiple testing correction was applied using the Benjamini–Hochberg method. The Python library XlsxReport (0.1.1) [7] was used to create formatted Excel files summarizing the results of the proteomics experiments.

##### *Data availability*

The mass spectrometry proteomics data have been deposited to the ProteomeXchange Consortium via the PRIDE [8] partner repository with the data set identifiers PXD063727, PXD063803, PXD063802, PXD063795, and PXD063731.

#### Phosphoproteomics

##### *Methanol/Chloroform precipitation and proteolytic digest*

Cortical P0 samples were extracted from n = 5 WT and homozygous litter-matched P0 animals of both the MAST1 L278del and the MAST1 G519S mouse line and flash-frozen for storage. The cortices were dissolved in 500 µL methanol and then protein was precipitated by adding 1 mL chloroform. After centrifugation (10 min, 10,000 g) the supernatant was removed. The pellets were lysed in a solution of 10 M urea and 50 mM HCl and then 1 M Triethylammoniumbicarbonate (TEAB) was added to a final concentration of 100 mM.

The lysates were supplemented with 5 mM DTT and incubated at 37°C for 1 h. The alkylation was performed by adding 10 mM Iodoacetamide and incubating for 30 min in the dark. The reaction was quenched by addition of 7.5 mM DTT and incubation at room temperature for 30 min. The samples were diluted to 6 M urea by addition of 100 mM TEAB and the proteins were digested with lysyl endopeptidase (Lys-C, Fujifilm Wako Pure Chemical Corporation) at an enzyme to protein ration of 1:50 for 3 h at 37°C. Subsequently, the samples were further diluted with 100 mM TEAB Buffer to 2 M urea, and trypsin (Trypsin Gold, Promega) was added at an enzyme to protein ratio of 1:50 and the samples were incubated overnight at 37°C.

The samples were acidified to a pH below 2 with 10% TFA and were desalted using C18 cartridges (Sep-Pak Vac 1cc (200 mg), Waters). Peptides were eluted with 2 x 600 µL 80% Acetonitrile (ACN) and 0.1% Formic Acid (FA), followed by freeze-drying.

*TMT labelling*

1.5 mg of lyophilized peptides were dissolved in 240 µL of 100 mM Triethylammonium bicarbonate. TMT10-plex labelling reagents (3 vials per label) were dissolved in 41 µL of anhydrous acetonitrile. 3 vials per label (2.4 mg of reagent) were added to the peptide samples and incubated for 1 h. After quenching with 5% hydroxylamine, the samples were mixed in several steps to achieve a correct mixing ratio of 1:1:1:1:1:1:1:1:1:1. (median of Top 500 proteins)

Following labelling, peptides were again freeze-dried, dissolved in TFA 0.1% and split in two aliquots: one containing 500 µg for measurement of the unmodified peptides and proteome quantification and one containing 13 mg for enrichment of phosphopeptides and phosphoproteome quantification. Both were desalted using C18 cartridges (Sep-Pak, Waters). Peptides were eluted with 70% ACN and 0.1% FA, followed by freeze-drying.

*Phosphopeptide enrichment*

Phosphopeptides were enriched using $TiO_2$ by first dissolving the dried peptide mixture in 4 mL of $TiO_2$ loading buffer (300 mg/mL lactic acid (Sigma-Aldrich), 80% ACN, 0.1% TFA) and incubating them with 120 mg of $TiO_2$ resin (Titansphere bulk media, 5 µm, GL Science) for 60 min at room temperature with over end rotation. For washing and elution the bead suspension was split and transferred to 8 Mobicol (MoBiTec) spin columns with a 10 µm pore filter at the outlet.

The $TiO_2$ beads in each spin column were washed with 8 x 500 µL of $TiO_2$ loading buffer, 8 x 500 µL of 80% ACN, 0.1% TFA, 4 x 500 µL of 1% ACN, 0.1% TFA, 4 x 500 µL of 1% ACN, 0.1% FA. Bound peptides were eluted from the resin by addition of 3 x 200 µL 0.3 M $NH_4OH$, 2 x 200 µL 0.7 M $NH_4OH$ and 2 x 150 µL 0.4 M $NH_4OH$ and 40% ACN, eluates were unified and freeze-dried.

*Fractionation of total peptides and enriched phosphopeptides by SCX*

The lyophilized peptide samples were dissolved in SCX buffer A (5 mM phosphate buffer pH 2.7, 15% ACN). 200 µg of the unmodified peptides and the total amount of the enriched phosphopeptides were loaded on the SCX column. The SCX fractionation was performed on an Ultimate system (Thermo Fisher Scientific) using a TSKgel SP-25W (ToSOH) column (5 µm particles, 1 mm i.d. x 300 mm) at a flow rate of 40 µL/min.

For the separation a ternary gradient was used, starting with 100% buffer A for 10 min, followed by a linear increase to 10% buffer B (5 mM phosphate buffer pH 2.7, 1 M NaCl, 15% ACN) and 50% buffer C (5 mM phosphate buffer pH 6.0, 15% ACN) in 15 min (for proteome) or 30 min (for phosphoproteome), to 25% buffer B and 50% buffer C in next 5 min (for proteome) or 10 min (for phosphoproteome), to 50% buffer B and 50% buffer C in next 5 min and an isocratic elution for further 15 min.

The flow-through was collected as a single fraction, along the gradient fractions were collected every minute and stored before analysis.

### *LC-MS/MS*

The nano HPLC system used was an UltiMate 3000 HPLC RSLC nano system (Thermo Scientific) coupled to a Q Exactive HF-X mass spectrometer (Thermo Scientific), equipped with a Proxeon nanospray source (Thermo Scientific). Peptides were loaded onto a trap column (Thermo Scientific, PepMap C18, 5 mm × 300 μm ID, 5 μm particles, 100 Å pore size) at a flow rate of 25 μL/min using 0.1% TFA as mobile phase. After 10 min, the trap column was switched in line with the analytical column (Thermo Scientific, PepMap C18, 500 mm × 75 μm ID, 2 μm, 100 Å). Peptides were eluted using a flow rate of 230 nL/min and a binary 120 min gradient. The two-step gradient started with the mobile phases: 98% A (water/formic acid, 99.9/0.1, v/v) and 2% B (water/acetonitrile/formic acid, 19.92/80/0.08, v/v/v) increased to 40% B over the next 120 min, followed by a gradient in 5 min to 95% B, stayed there for 5 min and decreased in 2 min back to 98% A and 2% B for equilibration at 30°C.

The Q Exactive HF-X mass spectrometer was operated in data-dependent mode, using a full scan (m/z range 380-1650, nominal resolution of 120,000, target value 3E6) followed by MS/MS scans of the 10 most abundant ions. MS/MS spectra were acquired using normalized collision energy of 35, isolation width of 0.7 m/z, resolution of 45,000, a target value of 1E5 and maximum fill time of 250 ms. For the detection of the TMT reporter ions a fixed first mass of 110 m/z was set for the MS/MS scans. Precursor ions selected for fragmentation (exclude charge state 1, 7, 8, >8) were put on a dynamic exclusion list for 30 s. Additionally, the minimum AGC target was set to 1E4 and intensity threshold was 4E4. The peptide match feature was set to preferred and the exclude isotopes feature was enabled.

*Data processing*

For peptide identification, the RAW-files were loaded into Proteome Discoverer (version 2.3.0.523, Thermo Scientific). All hereby created MS/MS spectra were searched using MSAmanda v2.3.0.12368, engine version 2.0.0.12368 [9]. The RAW-files were searched against the Uniprot reference database, using mouse as sub-organism (21,963 sequences; 11,719,000 residues). The following search parameters were used: Iodoacetamide derivative on cysteine, 11-plex tandem mass tag® (TMT) on lysine were set as fixed modifications, deamidation on asparagine and glutamine, oxidation on methionine, phosphorylation on serine, threonine and tyrosine as well as Carbamylation and 11-plex TMT on peptide-N-term were set as variable modifications. Monoisotopic masses were searched within unrestricted protein masses for tryptic enzymatic specificity. The peptide mass tolerance was set to ±5 ppm and the fragment mass tolerance to ±15 ppm. The maximal number of missed cleavages was set to 2. The result was filtered to 1% FDR on protein level using Percolator algorithm integrated in Thermo Proteome Discoverer. The localization of the modification sites within the peptides was performed with the tool ptmRS, which is based on phosphoRS [10]. Peptides were quantified based on Reporter Ion intensities extracted by the "Reporter Ions Quantifier"-node implemented in Proteome Discoverer. Proteins were quantified by summing unique peptides.

To distinguish regulated phospho-peptides that are altered due to regulation of the phosphorylation site from changes due to regulation of the underlying protein, PSMs were separated into those acquired from phospho-enriched samples and others acquired from the input before enrichment to capture changes on the proteome level. Therefore, proteins were quantified by summing unique and razor peptides detected and quantified in the input fractions. Resulting values were sum normalized across samples. Fractions of the phospho-enrichment were grouped and summarized on the peptide isoform level.

Subsequently, phosphopeptide regulations determined in the enriched fractions were corrected by regulation of the corresponding protein in the input fractions to correct for changes observed due to regulation on the proteome level. Statistical significance of differentially expressed phosphopeptides was determined using LIMMA [6] and FDR curves determined in Hein *et al* [12] were applied to extract significantly regulated phosphopeptides. MS/MS spectra assigned to the Tau peptide phosphorylated on residue S214 were validated manually. Most of them showed a mix of different isoforms, mainly of phosphorylated T212 and S214, while some spectra did not contain fragments that allowed for the unambiguous distinction between the possible isoforms.

##### *Data availability*

The phosphoproteomics data have been deposited to the ProteomeXchange Consortium via the PRIDE [10] partner repository with the data set identifier PXD065759.

#### Glutaraldehyde crosslinking

Purified MAST1$^{DK,}$ MAST1$^{D}$ p-S90, 14-3-3η, and co-purified MAST1$^{DK}$:14-3-3η complex was incubated either alone or in combination, at a final concentration of 20 µM, with 0.01% glutaraldehyde in buffer (50 mM HEPES pH 7.4, 150 mM NaCl, 1 mM TCEP), at room temperature for up to 30 min. The reaction was quenched with 20 mM Tris pH 7.5.

#### Generation of MAST1 G519S mouse model

Animals used in this study were housed at the IMP/IMBA animal facility under a 12:12 light-dark cycle. Food was provided ad-libitum. G519S MAST1 transgenic mice were generated by CRISPR/Cas9 editing through pronuclear microinjection using Cas9 mRNA, sgRNA, and a single-stranded DNA repair template [11]. See supplementary table 1 for gRNA and repair template sequences. The gRNA target sequence was cloned into PX458 plasmid as described previously [13] and sgRNA was expressed using the MEGAshortscript™ T7 Transcription kit (Thermo Fisher Scientific, AM1354). The ssDNA repair template was ordered from Integrated DNA Technologies (Alt-R™ HDR Donor Oligo). Injections were performed on BL6/CBA F1 zygotes. Successfully edited mice were identified using Sanger sequencing and backcrossed using C57BL/6J mice for at least 10 generations. All procedures were carried out in accordance with the legal requirements under an approved license (M58/006093/2011/14).

#### Solubility analysis of MAST1 mutants

MAST1$^{D}$ ΔS90WT, E194del, K276del, and L278del were transformed into BL21 Star *E. coli* and grown at 37°C to an OD of 0.7, before induction with 100 µM IPTG and overnight expression at 18°C. Cell pellets from 10 mL of expression culture were lysed in 2 mL of lysis buffer (50 mM HEPES pH 7.4, 150 mM NaCl, 1 mM TCEP, 2 mM benzamidine, 2 mM $MgCl_2$, 1x protease inhibitor cocktail (Sigma P8849), 0.2 kU Denarase, 2 mg lysozyme). Lysates were sonicated three times for 30 sec at 20% amplitude. A sample was taken for SDS-PAGE (whole cell lysate). The lysate was centrifuged at 38,397 g, 4°C, for 15 min, and a sample of the cleared lysate was taken for SDS-PAGE (supernatant). The pellet was resuspended in the same volume of buffer (50 mM HEPES pH 7.4, 150 mM NaCl, 1 mM TCEP, 0.25% CHAPS) by vigorously vortexing, and a sample was taken for SDS-PAGE (pellet).

#### Western blot

Mouse P0 cortical lysates were prepared using lysis buffer (20 mM Tris pH 7.5, 100 mM NaCl, 10% Glycerol, 1% Triton-X100) and 1:100 Halt™ Protease & Phosphatase Inhibitor Cocktail (Thermo Fisher Scientific, #78444). Brains were lysed by addition of 10 µL lysis buffer per mg of sample, addition of a Tungsten carbide bead and shaking using a TissueLyser II machine (Qiagen; 20 Hz, 2 x 60 s, RT) followed by further incubation on a rotating tube holder (60 min, 4°C). Protein concentrations were determined using a Pierce™ BCA Protein Assay Kit (Thermo Fisher Scientific, #23225). 30 µg of protein lysate was prepared in Laemmli loading buffer + 2% β-Mercaptoethanol and boiled for 5 min at 95°C. Gel electrophoresis was carried out using Novex™ WedgeWell™ 8-16% Tris-Glycine gel (Thermo Fisher Scientific, # XP08160BOX) and Tris-Glycine SDS running buffer (25 mM Tris, 240 mM glycine, 0.1% SDS, pH 8.3) for 2 h at 120 V. Blotting was done on activated PVDF membranes via wet transfer in transfer buffer (31 mM Tris, 240 mM Glycine, 0.025% SDS, 20% Methanol) for 16 h at 20 V. Membranes were blocked used 5% skimmed milk in TBS-T for 1 h @ RT. Incubation with primary antibody was performed overnight, 4°C in blocking solution. The following primary antibodies were used: Anti-MAST1 (ProteinTech, 13305-1-AP, 1:1000), Anti-MAST2 (SantaCruz, sc-377198, 1:300), Anti-MAST3 (Novus Biologicals, NBP1-82993, 1:500), Anti-GAPDH (Millipore, MAB374, 1:2000). Membranes were washed with TBST and incubated for 1 h, RT with the following secondary antibodies: Goat anti-mouse HRP (Abcam, ab6823, 1:5000), Goat anti-rabbit HRP (Abcam, ab6721, 1:10000). Chemiluminescent imaging was carried out using the ECL™ Western blotting reagent (Sigma Aldrich, # GERPN2109) on a Chemidoc-MP Bioimaging system (Bio-Rad). Quantification was performed using ImageJ.
